# RNAi and Ino80 complex control rate limiting translocation step that moves rDNA to eroding telomeres

**DOI:** 10.1101/2021.01.28.428704

**Authors:** Manasi S. Apte, Hirohisa Masuda, David Lee Wheeler, Julia Promisel Cooper

**Affiliations:** Telomere Biology Section, Laboratory of Biochemistry and Molecular Biology, National Cancer Institute, National Institutes of Health, Bethesda, MD 20892 USA; Department of Biochemistry, Cell and Molecular Biology, University of Tennessee, Knoxville, TN 37996 USA; Department of Biochemistry and Molecular Genetics, University of Colorado Anschutz Medical Campus, Aurora, CO 80045 USA

## Abstract

The discovery of HAATI^rDNA^, a mode of telomerase-negative survival in which canonical telomeres are replaced with ribosomal DNA (rDNA) repeats that acquire chromosome end-protection capability, raised crucial questions as to how rDNA tracts ‘jump’ to eroding, nonhomologous chromosome ends. Here we show that HAATI^rDNA^ formation is initiated and limited by a single translocation that juxtaposes rDNA from Chromosome (Chr) III onto subtelomeric elements (STE) on Chr I or II; this rare reaction requires the RNAi pathway and the Ino80 nucleosome remodeling complex (Ino80C), thus defining an unforeseen relationship between these two machineries. The unique STE-rDNA junction created by this initial translocation is efficiently copied to the remaining STE chromosome ends, without the need for RNAi or Ino80C, forming HAATI^rDNA^. Intriguingly, both the RNAi and Ino80C machineries contain a component that plays dual roles in HAATI subtype choice. Dcr1 of the RNAi pathway and Iec1 of the Ino80C both promote HAATI^rDNA^ formation as part of their respective canonical machineries, but both also inhibit formation of the exceedingly rare HAATI^STE^ (in which STE sequences mobilize throughout the genome and assume chromosome end protection capacity) in non-canonical, pathway-independent manners. This work provides a glimpse into a previously unrecognized crosstalk between RNAi and Ino80C in controlling unusual translocation reactions that establish telomere-free linear chromosome ends.

## Introduction

The linearity of eukaryotic chromosomes necessitates the assembly of telomeres, protective structures at chromosome termini that preserve genome integrity. Telomeres comprise arrays of short G-rich repeats culminating in a 3’ single-stranded (ss) overhang, and the shelterin complex, which includes double-strand (ds) telomere-binding proteins, ss overhang binding proteins, proteins that bridge these two, and telomeric noncoding RNAs (1–3). Shelterin integrates many activities that protect chromosomal termini from non-homologous end joining (NHEJ), extensive nucleolytic attack and homologous recombination (HR). Furthermore, shelterin engages and regulates the reverse transcriptase, telomerase, that replenishes the loss of telomeric repeats imposed by the end replication problem in each cell cycle (4).

While telomerase is expressed in the human germline, somatic cells inactivate its expression in early embryogenesis, resulting in progressive attrition of telomeres with each round of DNA replication. This process of ‘replicative senescence’ limits the lifespan of human somatic cells. In contrast, most (~85%) cancer cells reactivate telomerase to overcome this lifespan limitation. The remaining ~15% of tumor cells stabilize their chromosome ends and achieve unlimited proliferation by telomerase-independent strategies, collectively known as Alternative Lengthening of Telomeres (ALT). In the ALT cell lines characterized thus far, telomeres are heterogeneously maintained by break-induced replication (BIR), which is normally prevented at telomeres but promoted in ALT cells by a combination of replication stress and deregulated telomeric chromatin organization (5). In specific cancer types like liposarcomas, more than 50% of patient samples fail to show characteristics of any known telomere maintenance mechanisms (6,7), hinting at the presence of yet-unrecognized modes of linear chromosome maintenance deployed by cancer cells.

The fission yeast *Schizosaccharomyces pombe (S.pombe)* employs the classical telomere architecture to protect its chromosome ends. The ~300 base pairs (bp) of fission yeast telomere repeats (TTAC(A)GG(G_0-4_) engage a canonical shelterin complex comprising the dsDNA telomere binding protein Taz1 (ortholog of human TRF1and TRF2), ssDNA binding protein Pot1, and bridging proteins Rap1, Tpz1, Poz1 and Ccq1. Constitutive expression of both the reverse transcriptase (Trt1) and RNA (Ter1) subunits of telomerase obviates the end replication problem in unperturbed *S.pombe* cells. Deletion of genes encoding telomerase subunits leads to gradual telomere attrition, cell cycle arrest and death. Telomerase-negative survivors arise via three distinct mechanisms (8–10). The low chromosome number in *S.pombe* (three per haploid genome) allows formation of cells harboring three intra-chromosomal end fusions but no dicentric (and therefore lethal) inter-chromosomal fusions. Such circular chromosome-containing survivors, referred to hereafter as ‘circulars’ (O), are the most prevalent type of telomerase-minus survivors when cells are grown in non-competitive (single colony) conditions. Alternatively, telomeres can be heterogeneously maintained by continual homologous recombination, presumably via BIR between eroding telomeres, forming ‘linear’ (L) survivors.

The discovery of ‘HAATI’ (heterochromatin (HC) amplification-mediated and telomerase-independent), a third mode of telomerase-minus survival, established a new paradigm wherein telomere sequences *per se* are not essential for maintaining chromosome linearity (9). HAATI cells replace telomeres with blocks of non-telomeric HC that acquire the ability to prevent chromosome end fusions and maintain cell viability. HAATI emerges only under competitive growth conditions, as HAATI cells grow more rapidly than circulars and overtake the culture; in contrast, HAATI is exceedingly rare under non-competitive conditions. HAATI constitutes two sub-types. By far the most common is HAATI^rDNA^, in which ribosomal DNA (rDNA) repeats are copied from their wild-type (wt) loci on either end of Chromosome III (Chr III) to all chromosomal termini. In the exceedingly rare second subtype, HAATI^STE^, the rDNA remains at its original loci while the so-called sub-telomeric elements (STE) are amplified from the subterminal regions of Chr I and II to all chromosome termini as well as multiple internal loci. In both cases, terminal rDNA or STE repeats undergo continual expansion and contraction. However, the re-introduction of telomerase to HAATI cells (termed hereafter as HAATI + Trt1) results in the addition of canonical telomere repeats to HAATI chromosome ends, stabilizing the rearranged genomes.

Despite the absence of telomere repeats at HAATI^rDNA^ chromosome ends, the canonical shelterin factors Pot1 and Tpp1 are required for chromosome end-protection in these cells (9,11). Instead of recruiting Pot1/Tpp1 *via* telomere-specific DNA binding, HAATI^rDNA^ cells utilize rDNA-bound HC components including the histone methyltransferase Clr4, the HP1 protein Swi6, the histone deacetylase complex SHREC ((Snf2/HDAC-containing repressor complex), and a terminal ss rDNA overhang to recruit Pot1/Tpp1 and maintain HAATI^rDNA^ chromosome linearity. This Pot1/Tpp1 recruitment pathway is remarkably efficient whenever rDNA is present at subtelomeric regions, as HAATI^rDNA^ forms in 100% of telomerase-minus cells even under noncompetitive conditions if their genomes are ‘pre-rearranged’ (ie, in HAATI^rDNA^+Trt1 cells). Thus, the rarity of HAATI^rDNA^ formation stems solely from the step in which rDNA translocates to all chromosome ends. Intriguingly, we found that the RNAi pathway is essential for this translocation step, while RNAi is dispensable for HAATI^rDNA^ maintenance once rDNA has ‘jumped’ from Chr III to the termini of Chr I and II. As RNAi is generally prohibitive to the recombination associated with HC repetitive elements, the absolute dependence of rDNA jumping on RNAi was unexpected and remains to be mechanistically understood.

Whole-genome sequencing of several HAATI^rDNA^ survivors has underlined several additional facets of the translocations that underlie their formation. The preserved native transcriptional polarity of rDNA repeats after translocation rules out head-on end-joining as the translocation mechanism. Moreover, the sequences defining the sites of rDNA translocation localize to the rDNA intergenic spacer, a region in which origin firing, replication fork barrier (RFB) activity, RNA polymerase I (RNAPI) as well as Dcr1-restrained RNA polymerase II (RNAPII) transcription converge (12–14); the STE region is likewise a potentially unstable region subject to transcriptional repression (the telomere position effect, TPE) (15). These and other observations led us to postulate that upon telomere erosion, as TPE is lost and DNA damage responses are activated locally, subtelomeric regions sustain collisions between replication and transcription machineries, creating R-loops and/or unwound but unreplicated DNA. This instability may create the potential for template switching by either RNAPII or DNA polymerase, triggering the non-homologous translocations required for HAATI^rDNA^ formation.

Here we define the HAATI^rDNA^ translocation mechanism further. We find that the rarity of HAATI^rDNA^ formation stems from a single illegitimate recombination event that places rDNA at one STE chromosome end; this new STE-rDNA junction can then be copied efficiently onto the remaining chromosome ends. The RNAi pathway is critical only for the initiating step. We also identify a novel function for the Ino80 chromatin remodeling complex (Ino80C) in promoting this first translocation step. Intriguing parallels between the RNAi and Ino80 machineries emerge, as both machineries involve a protein that plays dual roles, promoting HAATI^rDNA^ formation in a pathway-dependent manner while inhibiting HAATI^STE^ formation in a pathway-independent manner. Hence, this work provides a glimpse into a previously unrecognized crosstalk between RNAi and Ino80C in controlling nonhomologous translocation reactions.

## Material and Methods

### Strains and media

*S. pombe* strains used in this study were derivatives of the standard laboratory strain 972 and are listed in Supplemental Table S1. Strains were grown at 32°C in standard rich medium (yeast extract with supplements-YE5S) unless indicated otherwise (16). Plasmid-containing strains were grown under selection for the appropriate marker. All tagging and gene deletions were constructed by one-step gene replacement (17), starting with the construction of heterozygous diploids except when noted. For reintroduction of telomerase, strains were transformed with p-kanMX-trt1^+^-myc (18).

### Generation of *trt1*Δ genomes with pre-existing STE-rDNA junctions on Chr I

HAATI^rDNA^+Trt1 cells with pre-arranged genomes were mated to *trt1+* cells in which Chr I was engineered to harbor genetic markers and fluorescent lacO/I arrays near either end of Chr I at A67:5479451 and sod2: 3185470 (19). Progeny harboring Chr I from a HAATI^rDNA^ cell and Chr II from a *wt* cell were selected, and chromosome arrangements were confirmed by pulsed field gel electrophoresis (PFGE; see below).

### Generating *trt1*Δ survivors

For competitive culturing in patches, equal volumes of *trt1*Δ cells were patched onto rich media plates and propagated for ~28 d by repatching equal volumes onto fresh plates every ~24 h. For noncompetitive culturing, *trt1*Δ offspring were streaked to single colonies iteratively for ~28 d.

### Dilution assays

Cells were grown in liquid culture to log phase, and culture density measured using a hemocytometer and adjusted to 1 × 10^7^ cells per milliliter. Five-fold serial dilutions were pipetted in a 96-well microtiter plate with repeated agitation. Diluted cells were stamped onto plates using a metal stamper (which transfers ~5 μL). For determining MMS sensitivity, freshly prepared YES agar containing either DMSO control or 0.001%, 0.003% and 0.006% v/v MMS were used. MMS sensitivity was determined after incubation at 32°C for 3-4 days. All experimental strains were compared against WT, Circular (O) and pre-existing HAATI strains as controls, As compared to WT, O survivors are extremely MMS sensitive while HAATI survivors are relatively MMS resistant.

### DNA isolation and Southern blotting

DNA isolation and Southern blot analysis were performed as described previously (20).

### PFGE

PFGE of whole chromosomes was performed as described previously (9) with the following modifications: Agarose plugs were loaded onto 0.8% agarose gels in 1X TAE (40 mM Trisacetate buffer, 2 mM Na2EDTA at pH 8.3). PFGE was performed on a BioRad CHEF DR-III system in 1X TAE at 14°C using the following program: step 1, 31 h at 2 V/cm, 96° angle, and 1200-sec switch time; step 2, 31 h at 2 V/cm, 100° angle, and 1500-sec switch time; and step 3, 31 h at 2 V/cm, 106° angle, and 1800-sec switch time. After electrophoresis, DNA was visualized by ethidium bromide staining, and gels were processed for Southern blot analysis using the STE1 probe (8) or the rDNA probe (21).

PFGE of NotI-digested chromosomes was performed as described previously (9) with the following modifications: NotI-digested agarose plugs were loaded onto 1% agarose gels in 0.5X TBE (1 × TBE: 89 mM Tris-borate, 2 mM EDTA). PFGE was performed on a Bio-Rad CHEF DR-III system in 0.5X TBE at 14°C using the following program: 27 h at 6 V/cm, 120° angle, and 60- to 120-sec switch time. After electrophoresis, DNA was visualized by ethidium bromide staining, and gels were subjected to Southern blot analysis using LMIC probes (22) or the STE1 probes.

### Whole Genome Sequencing and analysis

Whole genome sequencing of single HAATI^rDNA^ clones and analysis were performed as described previously (11).

## Results

### Whole genome sequencing suggests two-step model for translocation of rDNA to all chromosome ends

Whole genome sequencing of several *trt1*Δ HAATI^rDNA^ genomes had shown that they differ from *wt* genomes only at their chromosome termini, and that translocated rDNA sequences preserve their polarity (rDNA transcription towards chromosome termini), suggesting that HAATI^rDNA^ arise via a DNA- or RNA-polymerase template-switching event that creates novel a STE-rDNA junction at the interface of each translocation. Intriguingly, while the precise location and sequence of these junctions varies between clones, our initial sequencing experiments suggested that within a single HAATI^rDNA^ clone, only one specific STE-rDNA junction could be found; i.e., every chromosome end in a given HAATI^rDNA^ cell appeared to harbor precisely the same STE-rDNA junction. To determine whether we could be missing additional junctions due to insufficient genome coverage, whole genome sequencing was performed at 300X coverage of the *S. pombe* genome (Figure 1A). Remarkably, even at this level of coverage, each clone contains only a single STE-rDNA junction. Thus, we hypothesize that the translocation placing rDNA from Chr III onto the STE-containing termini of Chr I and II occurs via two steps (Figure 1B). The first step comprises a single, rare translocation of rDNA to an eroding STE region. This generates a unique STE-rDNA junction for the first time within the *trt1*Δ genome. In the second step, this STE-rDNA junction is copied onto the remaining STE chromosome ends by ‘standard’ BIR. This copying of a single STE-rDNA junction results in identical STE-rDNA junctions at all of the chromosomal termini within single HAATI clone.

**Figure 1.**
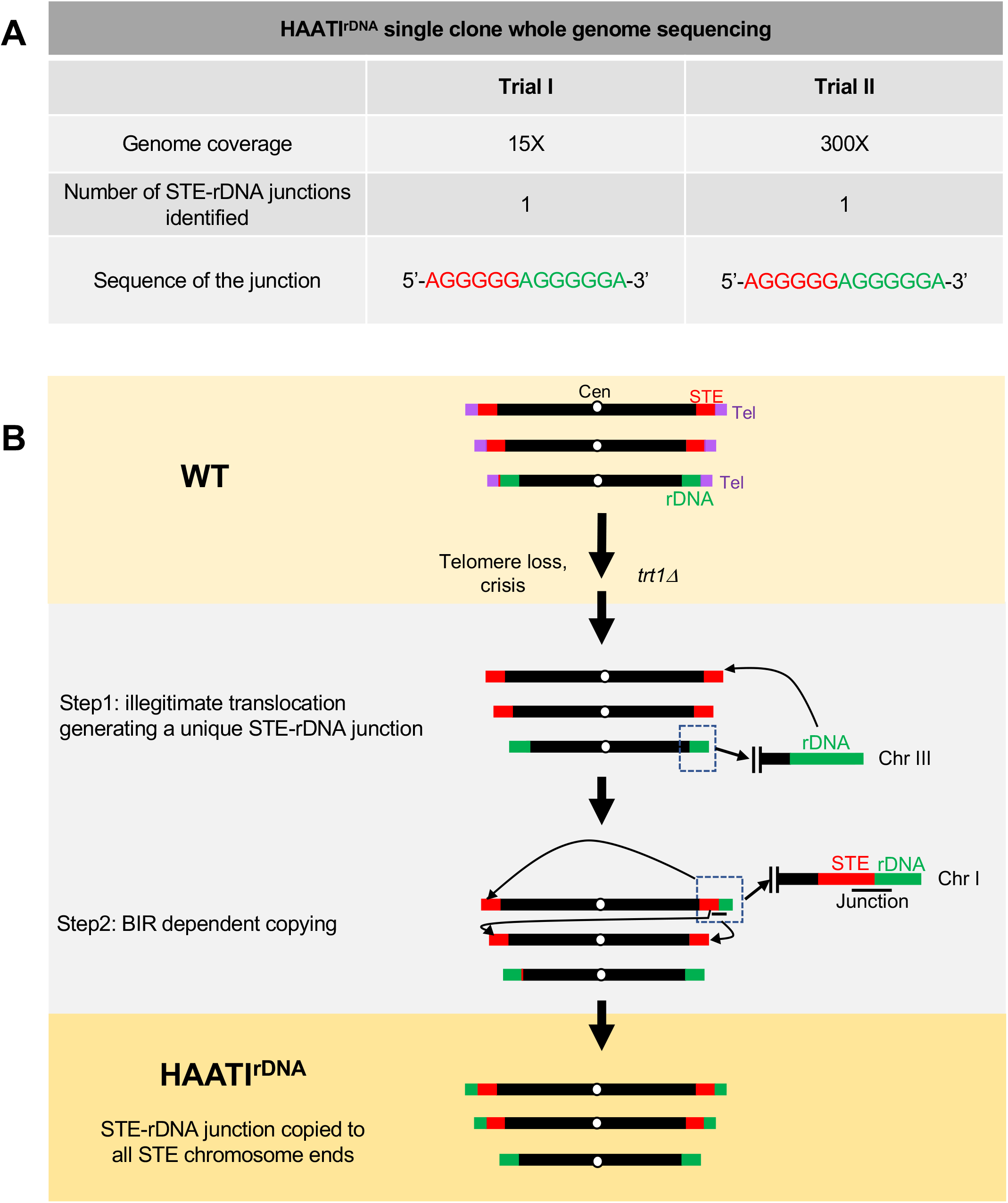
Whole genome sequencing suggests two-step model for HAATI^rDNA^ translocation. A. Comparative analysis of whole genome Illumina sequencing of a single HAATI^rDNA^ clone. Regardless of the genome coverage (15X vs 300X), only a single STE-rDNA junction is identified. B. Proposed two-step model for rDNA translocation. Linear chromosomes of S.pombe genomes are represented with telomeres (purple) flanked by STE (STE, red) on Chr I and II, and sub-terminally positioned rDNA on Chr III (green). Upon telomerase loss, HAATI^rDNA^ is proposed to arise via two steps: First, a unique STE-rDNA junction (black solid line) arises wherein a single illegitimate translocation places rDNA from Chr III on a STE tracts of Chr I or II. Second, the STE-rDNA junction is copied to the remaining STE-containing chromosomal ends, presumably via BIR.

### Single, rare, illegitimate translocation drives HAATI formation

Our previous work showed that the translocation of rDNA from Chr III to the ends of Chr I and II is rate-limiting for HAATI^rDNA^ formation. The foregoing two-step model in which rDNA is first placed at one nonhomologous site and then copied *via* BIR to the remaining STE chromosome ends further predicts that the rarity of HAATI^rDNA^ formation stems from the first single translocation. If this model is correct, provision of one (or in this case two) pre-existing STE-rDNA junctions should be sufficient to obviate the first, and rate-limiting step. To test this, we sought to create *trt1* Δ genomes with pre-existing STE-rDNA junctions on only one chromosome (Figure 2A-B). HAATI^rDNA^+Trt1 cells containing a 5’-STE-AGGGGG/AGGGGGA-rDNA-3’ junction, the most prevalent junction among clones thus far sequenced, were mated with *trt1+* cells in which Chr I was engineered to harbor genetic markers and fluorescent lacO/I arrays near either end of Chr I (at *sod2+,* 80 kb from Chr IL and A67, 100 kb from Chr IR). These features allow determination of whether Chr I has been inherited from the HAATI^rDNA^ parent or the *trt1+* parent. Progeny harboring Chr I from a HAATI^rDNA^ cell and Chr II from a *wt* cell (note that Chr III is flanked by rDNA in both *wt* and HAATI cells) were selected, and the presence of the STE-rDNA junctions on Chr I confirmed by PFGE (Figure 2C).

**Figure 2.**
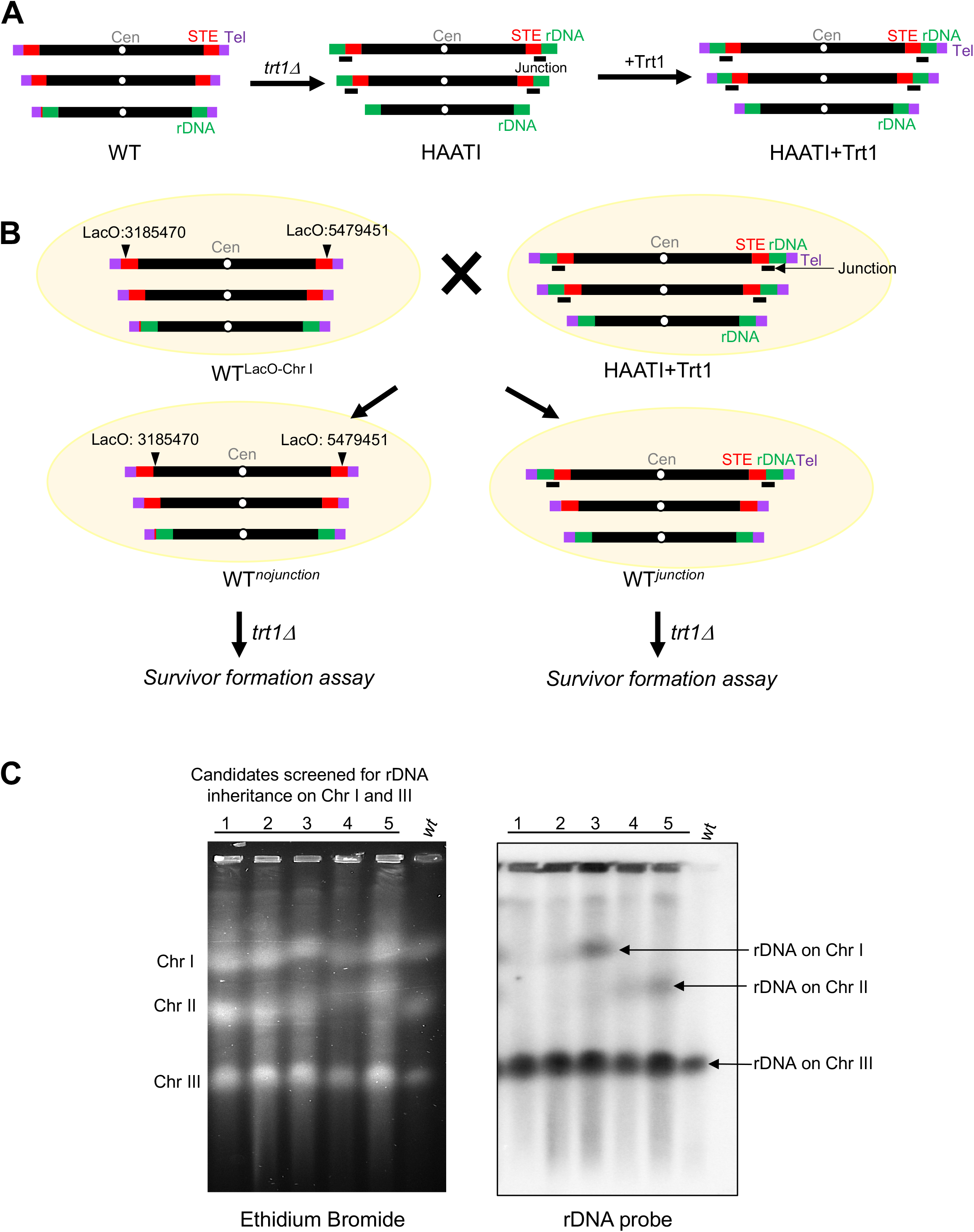
Generation of *S.pombe* genome with a pre-existing STE-rDNA junction on Chr I. A. When plasmid-borne Trt1 is introduced into HAATI^rDNA^ cells, telomeres are added and the rearranged genomes (with rDNA at all subtermini) are stabilized; these HAATI^rDNA^+Trt1 genomes are ‘pre-rearranged’. B. Generation of cells harboring a single HAATI^rDNA^+Trt1 chromosome. *wt* cells containing LacO repeats integrated at the A67: 5479451 and Sod2: 3185470 loci on Chr I were mated with HAATI^rDNA^+Trt1 cells. Offspring lacking or containing LacO repeats were selected via nearby selectable markers. Offspring lacking the LacO repeats have inherited Chr I from the HAATI^rDNA^+Trt1 parent (with STE-rDNA junctions near either chromosome end), and were further screened for the absence of rDNA on Chr II (C). C. Whole chromosome PFGE reveals which chromosomes have been inherited from HAATI^rDNA^+Trt1 parent. Several *trt1*Δ clones (1–5) that lack or contain LacO repeats on Chr I are shown. Ethidium bromide staining shows the presence of all three chromosomes (left); Southern blot analysis with an rDNA probe (right) shows that in *wt* genomes (WT lane), rDNA is restricted to Chr III. Based on rDNA signal on Chr I or II, *trt1Δ^unction^* (carrying Chr I from HAATI+*Trt1* parent) and *trt1 Δ^nojunction^* (carrying Chr I without STE-rDNA junction) were selected for subsequent experiments.

*trt1*Δ cell populations lacking (Figure 3A, *trt1 Δ^nojunction^*) or carrying a STE-rDNA junctions (Figure 3B, *trt1 Δ^junction^)* were tested for frequency of HAATI^rDNA^ formation under competitive and non-competitive growth conditions. In otherwise *wt trt1*Δ genomes, virtually 100% of survivors raised under competitive growth conditions (liquid or serial patching) use the HAATI^rDNA^ strategy while under non-competitive growth conditions (repetitive single colony streaking), circular survivors dominate. Survivor type was initially assessed by testing MMS sensitivity (9), as severe MMS sensitivity, a hallmark of ‘circular’ survivors, is averted by the HAATI survival mode, whether cells are HAATI^rDNA^ or HAATI^STE^. As expected, 10 of 10 *trt1 Δ^nojunction^* genomes formed HAATI under competitive conditions, but none form HAATI under non-competitive conditions (Figure 3A, top). However, *trt1*Δ genomes with pre-existing STE-rDNA junctions only on Chr I *(trt1 Δ^junction^*) generate HAATI survivors in 100% of cases, regardless of growth conditions (Figure 3A, bottom).

**Figure 3.**
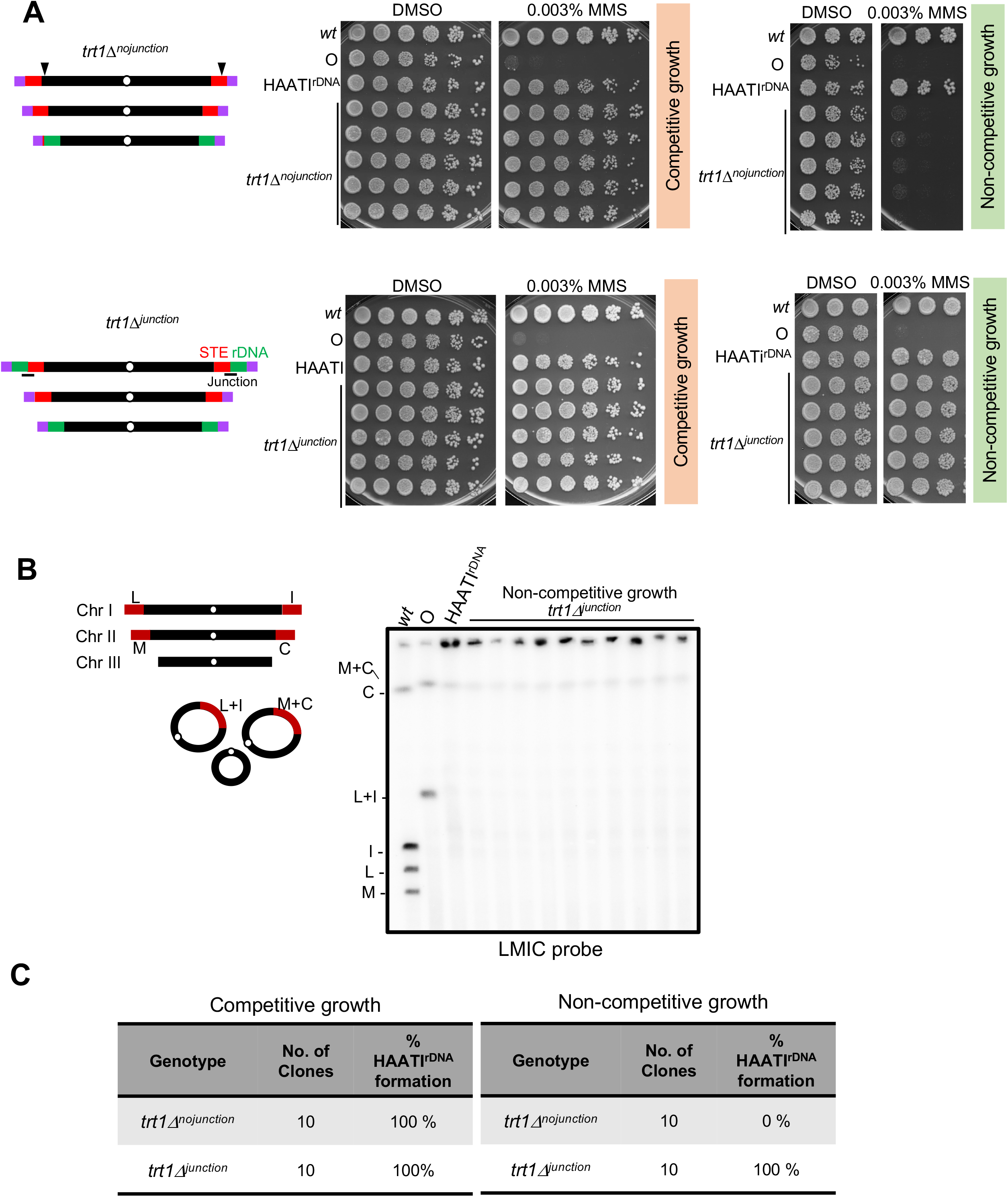
A pre-existing STE-rDNA junction guarantees HAATI^rDNA^ formation irrespective of growth conditions. A. Left: Schematic of *trt1 Δ^nojunction^* genome (top) or the *trt1Δ^junction^* genome (bottom). Right: Five-fold serial dilutions spotted on rich media with or without MMS. *trt1*Δ populations lacking STE-rDNA junctions form HAATI^rDNA^ under competitive conditions and circular survivors under noncompetitive conditions. *trt1 Δ^junction^* survivors show MMS resistance indicative of HAATI formation when raised under either condition. B. Validation of genome arrangements in *trt1 Δ^junction^* survivors raised by non-competitive growth by PFGE. NotI digestion of *wt* chromosomes releases four terminal fragments (L, M, I, and C) from Chr I and II; in circulars, these are replaced by fusion fragments L+I and C+M, while in HAATI cells, the vast majority of NotI fragments are retained in the well. All *trt1Δ^junction^* survivors are HAATI. C. Table summarizing the effect of a pre-existing junction on HAATI^rDNA^ formation. Percentage of HAATI^rDNA^ formation was scored based on MMS resistance, PFGE pattern and the absence of STE amplification (data not shown).

Survivors were further investigated by PFGE analysis of chromosome organization. Digestion of linear chromosomes with the rare cutter NotI releases four terminal fragments – (L, I, M and C; a probe cocktail with all four is referred to as LMIC) from linear Chr I and II, while circular chromosomes yield fused terminal fragments (L+I and M+C) (Figure 3B, left); in contrast, the terminal fragments of HAATI chromosomes fail to enter gels (presumably due to the continual presence of branched recombination intermediates), resulting in retention of LMIC hybridization in the well (Jain et al. 2010). As expected, all the MMS resistant survivors arising in *trt1*Δ^junction^ genomes, whether raised under competitive or non-competitive conditions, retained LMIC hybridization signal in the well, confirming their HAATI status (Figure 3B, right, 3C). We further validated that all were HAATI^rDNA^ survivors, as STE probe hybridization is absent (data not shown). Thus, the rarity of HAATI stems from the rarity of a first translocation to form one STE-rDNA junction, after which the junction is efficiently and accurately copied to additional STE chromosome ends, presumably by BIR. Bypassing this step allows HAATI to dominate even under non-competitive growth conditions.

### RNAi pathway is essential only for driving the first translocation step

The RNAi pathway is essential for the rDNA jumping that places rDNA tracts at all HAATI^rDNA^ chromosomal termini (11). To determine whether RNAi is required for the first, rate limiting translocation to form a single STE-rDNA junction or rather for the subsequent copying of this junction to remaining STE regions, we generated *trt1*Δ cells harboring or lacking *dcr1+* with pre-existing STE-rDNA junctions on Chr I (as described in Figure 2A-B). In otherwise *wt trt1*Δ genomes with no STE-rDNA junctions, loss of Dcr1 completely abolishes HAATI^rDNA^ formation (Figure 4A-B) (11). Remarkably, however, in *trt1*Δ genomes with pre-existing STE-rDNA junctions on Chr I, Dcr1 is dispensable for HAATI^rDNA^ formation (Figure 4C-E). While 0 of 10 *dcr1Δ trt1*Δ genomes lacking a junction form HAATI^rDNA^, 100% of *dcr1Δ trt1*Δ genomes with STE-rDNA junctions on Chr I do so, in all growth conditions (Figure 4F). Moreover, provision of STE-rDNA junctions on Chr I also bypasses the requirement of Ago1 for HAATI^rDNA^ formation (Figure 4F, Figure S1). Hence, the RNAi pathway is essential specifically for initial translocation that places rDNA next to STE tracts, and dispensable for subsequent copying of this junction to additional STE chromosome ends.

**Figure 4.**
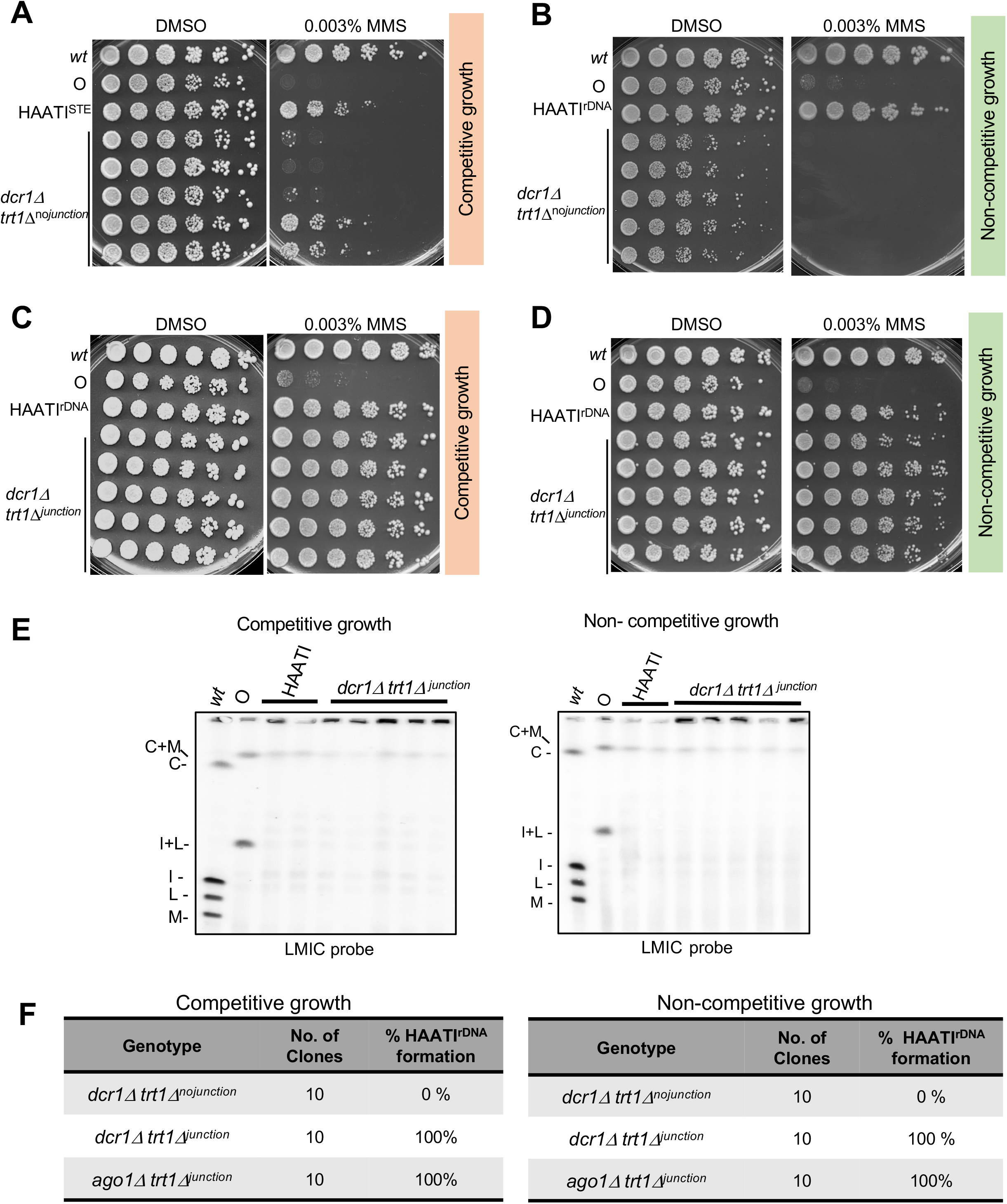
RNAi pathway is essential only for the initial translocation step. A. *dcr1Δ trt1 Δ^nojunction^* survivors propagated under competitive growth conditions and spotted in 5-fold serial dilutions yield MMS sensitive (upper 3 rows) and resistant (lower 2 rows) clones. The MMS resistant clones are presumably HAATI^STE^. B. *dcr1Δ trt1 Δ^nojunction^* survivors when propagated under non-competitive growth conditions showed extreme MMS sensitivity, indicating circular survivors. C. *dcr1Δ trt1 Δ^junction^* survivors propagated under competitive growth show MMS resistance indicative of HAATI formation. D. *dcr1 trt1 Δ^junction^* survivors propagated under non-competitive growth show MMS resistance indicative of HAATI formation. E. Determination of genome arrangements in *dcr1Δ trt1Δ^junction^* survivors by PFGE of Not1 digested chromosomes. Retention of hybridization signal in the well for all survivors indicates formation of HAATI. F. Table summarizing the dispensibility of RNAi factors for HAATI^rDNA^ formation when a pre-existing STE-rDNA junction is present. Percentage of HAATI^rDNA^ formation is based on MMS resistance, retention of PFGE signal and absence of STE amplification (data not shown).

### Telomere-proximal STE are preferred sequences for nonhomologous translocation, while knob regions are inhibitory

As all *wt* telomeres are flanked by repetitive regions, either STE or rDNA, we wondered whether rDNA always jumps to STE during HAATI^rDNA^ formation because of the special properties of STE sequences or because of their subterminal location. To approach this question, we asked whether rDNA can jump onto subtelomeric genomic sequences other than STE tracts, utilizing *STE*Δ strains in which the STE tracts on all chromosomal termini have been almost entirely replaced with *ura4+* or *his7+* selection markers (Figure 5A) (23). These *STE*Δ strains show growth rates and sporulation efficiencies resembling those of *wt* strains. We generated *trt1+* deletions in these *STE*Δ strains by sporulating *trt1^+/Δ^ STE^ΔΔ^* diploids and compared HAATI formation frequency under competitive growth conditions. Interestingly, the survivors arising in the absence of STE tracts are all circulars resulting from intra-/inter-chormosomal fusions, suggesting that even under competitive conditions where it normally dominates, HAATI^rDNA^ cannot form in the *STE*Δ setting (Figure 5B-C).

**Figure 5.**
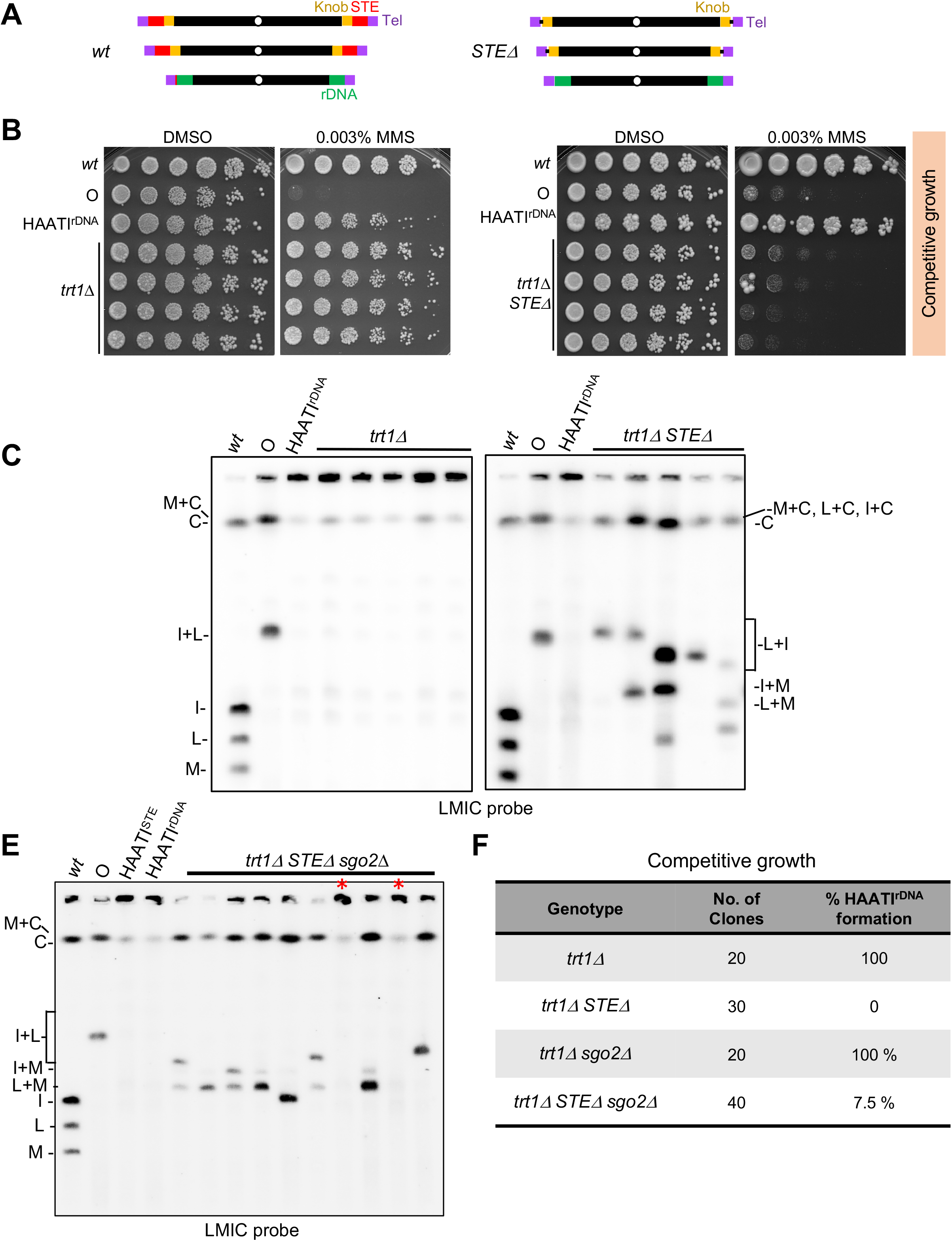
STE are preferred ‘acceptor sequences’ for HAATI^rDNA^ translocation. A. *wt S.pombe* genomes contain knob regions (yellow), whose chromatin marks and repressive properties that depend on Sgo2 recruitment, just proximal to STE regions. STExΔ strains lack virtually all STE tracts, placing the knob regions in close proximity to the canonical telomeric repeats. B. *trt1*Δ survivors obtained under competitive growth conditions and spotted in 5-fold serial dilution show MMS resistance indicating HAATI formation. *trt1Δ STE*Δ survivors obtained under the same conditions show extreme MMS sensitivity indicating chromosome circularization. C. Genome arrangements of survivors. (Left) Retention of hybridization signal in the well for all survivors indicates HAATI formation. (Right) *trt1Δ STE*Δ survivors form inter- and intra-chromosomal fusions indicated by various size circularization products sometimes accompanied with chromosome length changes and/or dichromosome circles D. PFGE of NotI digested chromosomes of *trt1Δ STEΔ sgo2*Δ survivors. 2 out of 10 tested survivors (indicated by red asterisks) show retention of PFGE signal in the well, suggesting HAATI formation. E. Table summarizing the role of STE sequences and the Sgo2 bound knob on HAATI^rDNA^ formation. Percentage of HAATI^rDNA^ formation is based on MMS resistance, retention of PFGE signal and absence of STE amplification (data not shown).

While the foregoing experiments suggest that STE tracts are favored ‘acceptor’ regions for the translocations that lead to HAATI^rDNA^, it is also conceivable that the newly positioned telomere-proximal DNA in *STE*Δ genomes may assemble a chromatin environment that inhibits translocation. Indeed, the telomere-proximal region in *STE*Δ genomes (ie, the STE-proximal region in *wt* cells) is unusual, forming a domain that stains densely with DAPI and is termed the ‘knob’. The knob is a transcriptionally repressive domain, but unlike canonical heterochromatin which harbors histone H3K9-Me, the knob lacks H3K9Me and instead harbors deacetylated and methylated histone H3K36 (24,25). This modification, as well as the DAPI-dense staining of the knob, depend on the Shugoshin Sgo2 (25). We sporulated a *trt1Δ/+ STEΔ/Δ sgo2Δ/+* diploid to ask whether Sgo2 influences HAATI^rDNA^ formation. In otherwise *wt* genomes, Sgo2 is dispensable for HAATI^rDNA^ formation and maintenance (Figure S2A-B). Remarkably, however, loss of Sgo2 in *trt1Δ STE*Δ genomes results in a low frequency of HAATI^rDNA^ formation (7.5%, 3 out of 40 clones across three independent experiments), which contrasts with the complete absence of HAATI^rDNA^ formation in *trt1ΔSTE*Δ cells with intact Sgo2 (Figure 5E-F). This suggests a scenario in which the knob region is refractory to rDNA translocation, but the loss of Sgo2 renders this region weakly permissible for the illegitimate translocations that result in HAATI^rDNA^.

We wondered if the low frequency of HAATI^rDNA^ formation in *trt1Δ STEΔ sgo2*Δ genomes might stem from the residual Sgo2 function provided by its heterozygosity in the parental diploid. Conceivably, the Sgo2+ copy in this diploid provides knob-like properties that persist via epigenetic means in the offspring. To test this possibility, we attempted to generate *trt1+/*Δ diploids with both copies of *sgo2+* deleted. This led to abnormally shaped meiocytes evincing significant chromosome missegregation and asci with variable numbers of spores (2-8 per ascus), consistent with previous reports of Sgo2’s role in regulating meiosis (26). Due to the severity of this phenotype, we were unable to utilize *sgo2Δ/*Δ diploids to assay HAATI formation. Collectively, we surmise that the knob region is inhibitory to HAATI^rDNA^ translocations and speculate that knob function can persist in the offspring of *sgo2+/*Δ diploids, limiting the level of relief of this inhibition.

### Reb1 regulates HAATI^rDNA^ survival

The sequences within the rDNA intergenic spacer region that translocate during HAATI^rDNA^ formation bind Reb1, which provides a replication barrier, enforcing unidirectional replication fork passage in this region (13,27). Thus, we asked whether Reb1 is critical for HAATI survival. Deletion of *reb1+* in pre-formed HAATI^rDNA^ cells results in chromosome circularization, as evinced by extreme MMS sensitivity (data not shown) as well as PFGE signals indicating the fusion fragments I+L and M+C (Figure S3A). This effect is specific to HAATI^rDNA^, as *reb1+* deletion in pre-formed HAATI^STE^ had no effect on chromosome linearity. Furthermore, 100% of *reb1Δ trt1*Δ survivors raised under competitive conditions fail to form HAATI and instead formed ‘circular’ survivors (Figure S3B-C). Thus, Reb1 is crucial for HAATI^rDNA^ survival.

### Ino80 complex is essential for HAATI^rDNA^ formation

As the termini of HAATI^rDNA^ genomes must both maintain a HC structure that affords Pot1 recruitment, and transiently harbor chromatin with a sufficiently ‘open’ structure to allow the translocations that form STE-rDNA junctions, we sought to address how are these disparate chromatin environments are regulated. In this vein, we explored the role of an evolutionarily conserved ATP dependent chromatin remodeler, the Ino80 complex (Ino80C)(28–30) in HAATI^rDNA^ formation. We first probed the role of Arp8, a conserved subunit of the Ino80C, required for the nucleosome remodeling activity of Ino80C. We generated *trt1*Δ cells carrying or lacking Arp8, propagated them under competitive conditions that normally confer HAATI^rDNA^ formation, and analyzed survivor type by MMS sensitivity assay (data not shown) and PFGE. Loss of Arp8 completely abolishes HAATI^rDNA^ formation, yielding exclusively linear survivors (Figure 6A). Thus, Arp8 is required for HAATI^rDNA^ establishment and may be required for circular formation. Deletion of *arp8+* from pre-formed HAATI^rDNA^ cells leads to a reduced growth rate and enhanced MMS sensitivity (Figure S4A), but examination of chromosome structure indicates that these cells retain the linear HAATI^rDNA^ chromosome structure (Figure S4B).

**Figure 6.**
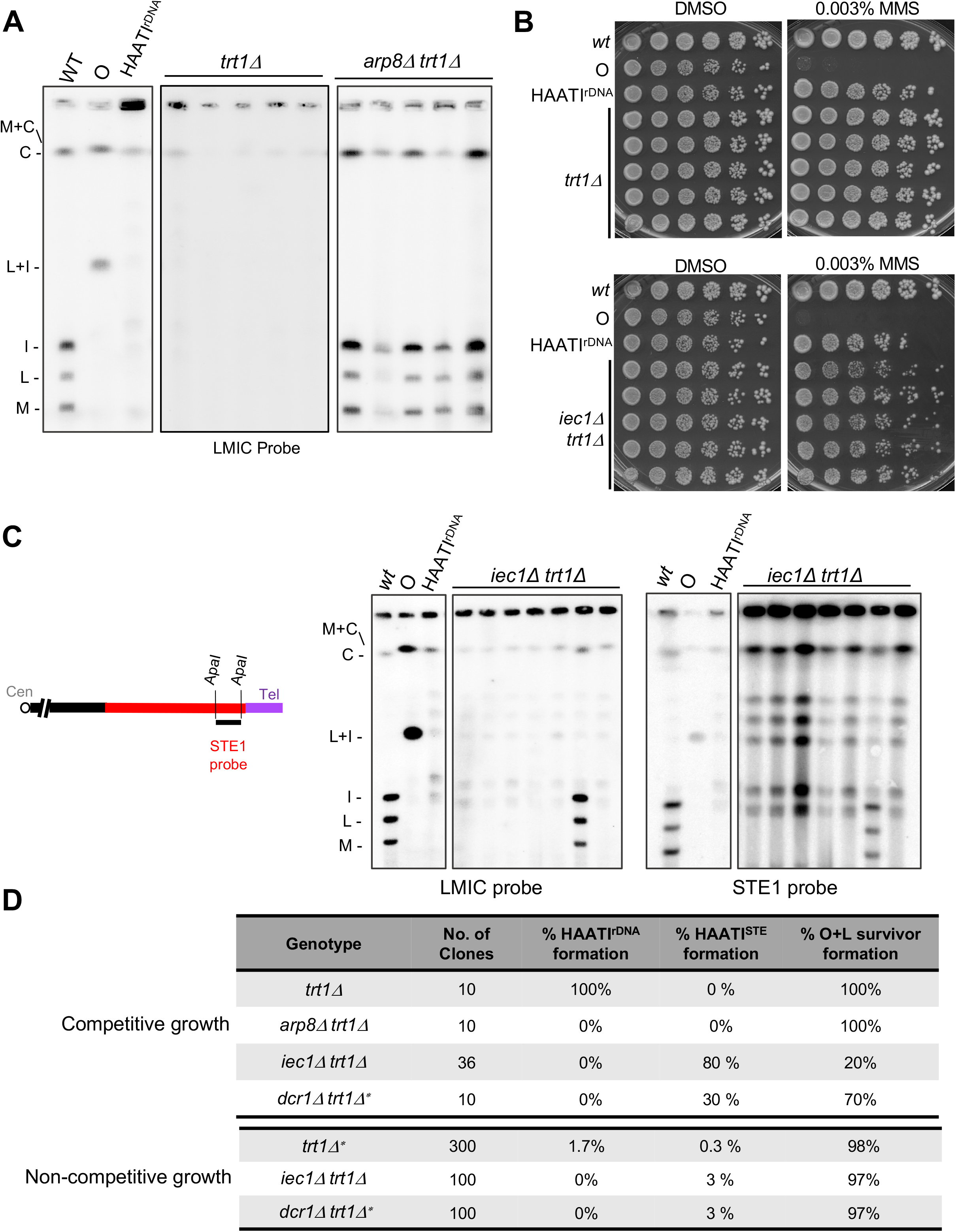
Ino80C is essential for HAATI^rDNA^ formation while Iec1, an Ino80C member, phenocopies Dcr1 in its role in HAATI subtype choice. A. progeny of heterozygous *trt1Δ/trt1+* diploids carrying or lacking *arp8^+^* were cultured under competitive conditions. HAATI survivors are absent upon loss of Arp8; instead, linear survivors form as indicated by PFGE pattern. B. Heterozygous *trt1Δ/trt1+* diploids carrying or lacking *iec1^+^* were sporulated, and the indicated progeny cultured under competitive conditions and spotted in 5-fold serial dilution. All *iec1Δ trt1*Δ survivors show MMS resistance indicating HAATI formation. C. (Left) Schematic shows location of STE1 probe (black line) in STE region (red). (Right) Representative PFGE of NotI-digested chromosomes from *iec1Δ trt1*Δ survivors. Most *iec1Δ trt1*Δ survivors are HAATI as indicated by retention of signal in the well probed for LMIC. The membrane was stripped and re-probed for STE1, which hybridizes strongly with all NotI restriction fragments, indicating HAATI^STE^ formation. D. Tables summarizing the essential role for Arp8 and Iec1 in HAATI^rDNA^ formation as well as Iec1’s role in blocking HAATI^STE^ formation independent of Ino80C, phenocopying Dcr1. Analysis of survivor formation is shown under competitive conditions (top) and noncompetitive conditions (bottom); asterisk marks published results from (11) shown here for comparison.

### Iec1 phenocopies Dcr1 in its roles in HAATI formation

To determine whether the requirement for Arp8 for HAATI^rDNA^ formation reflects a role for the Ino80C, we probed a second Ino80C member, Iec1. Two of ten pre-formed HAATI^rDNA^ clones that undergo loss of Iec1 circularize their chromosomes (Figure S4C); hence, Iec1 is partially required for the linear chromosome maintenance of HAATI^rDNA^. Along with the reduced growth rate seen upon loss of Arp8 in pre-formed HAATI^rDNA^, this suggests an important but not essential role for Ino80C in HAATI^rDNA^ maintenance.

In contrast, Iec1 has a dramatic impact on HAATI formation. When *iec1Δ trt1*Δ survivors are raised under competitive growth conditions, the majority of populations (80%) yielded an MMS resistance level characteristic of HAATI (Figure 6B). Surprisingly, however, PFGE analysis reveals STE1 probe hybridization to all of the internally located NotI fragments from all of these *iec1Δ trt1*Δ i solates (Figure 6C). This pattern is diagnostic of a rare subtype of HAATI, HAATI^STE^. Of 36 survivors analyzed by PFGE in three independent HAATI formation experiments, we never isolated an *iec1Δ trt1*Δ HAATI survivor lacking STE amplification. Hence, loss of Iec1 completely abolishes HAATI^rDNA^ formation while promoting HAATI^STE^ formation. The elevated level of HAATI^STE^ formation in the absence of Iec1 suggests that the rarity of HAATI^STE^ stems from an active role for Iec1 in inhibiting its establishment. To explore this idea, we assessed the *iec1Δ trt1*Δ survivor formation under non-competitive conditions, under which 3% of *iec1Δ trt1*Δ colonies are HAATI^STE^ while 97% are circulars (Figure 6D). This contrasts sharply with the 0.3% HAATI^STE^ formation rate seen in the presence of Iec1 (11), and confirms that upon telomerase loss, Iec1 actively inhibits HAATI^STE^ establishment. As HAATI^STE^ survivors were never obtained from *arp8Δtrt1*Δ populations even under competitive growth conditions, Arp8 does not share Iec1’s role in blocking STE mobilization.

The foregoing results show that Iec1 inhibits the establishment of HAATI^STE^ independently of canonical Ino80C. At the same time, Iec1 acts as part of the canonical Ino80C to promote HAATI^rDNA^ establishment. This role for Iec1 in survivor pathway choice is reminiscent of that of Dcr1 (11), which acts independently to suppress HAATI^STE^ formation, while acting in concert with the RNAi pathway to promote HAATI^rDNA^ formation (Figure 6D). These parallels between the involvement of Dcr1 and Iec1 in HAATI^rDNA^ establishment and HAATI^STE^ inhibition prompted us to explore a potential crosstalk between these two regulators. We first tested tandem deletions of *dcr1+* and *iec1+* in already-formed HAATI^rDNA^ and found no effect on HAATI^rDNA^ chromosome maintenance (Figure S5A). Next, we generated *trt1*Δ cells lacking both Iec1 and Dcr1, and propagated the survivors under competitive conditions. Remarkably, *dcr1Δiec1Δ trt1*Δ survivors failed to form HAATI (Figure S5B-C). Thus, while Dcr1 inhibits HAATI^STE^ formation in otherwise *wt trt1*Δ survivors, it is required for HAATI^STE^ formation in *iec1Δ trt1*Δ cells; conversely, Iec1 inhibits HAATI^STE^ formation in the presence of Dcr1, yet is required for HAATI^STE^ formation in Dcr1’s absence.

### Ino8C is essential only for the first translocation step

We sought to understand whether the Ino80C, like the RNAi pathway, is required solely for the rare STE-rDNA junction-forming step of HAATI^rDNA^ establishment. Deletions of genes *(arp8*Δ and *iecΔ)* encoding Ino80C components were generated in *trt1*Δ genomes containing or lacking pre-existing STE-rDNA junctions on Chr I, and survivors were raised under competitive and non-competitive growth conditions. Remarkably, while *iec1Δ trt1*Δ or *arp8Δ trt1*Δ never form HAATI^rDNA^ in the absence of a pre-existing STE-rDNA junction (Figure 6), HAATI^rDNA^ emerged in 100% of cases when a pre-existing STE-rDNA junction was present, regardless of growth conditions in both settings (Figure 7A-D). Hence, the role for the Ino80C in HAATI^rDNA^ establishment lies solely within this crucial first translocation to form a unique STE-rDNA junction.

**Figure 7:**
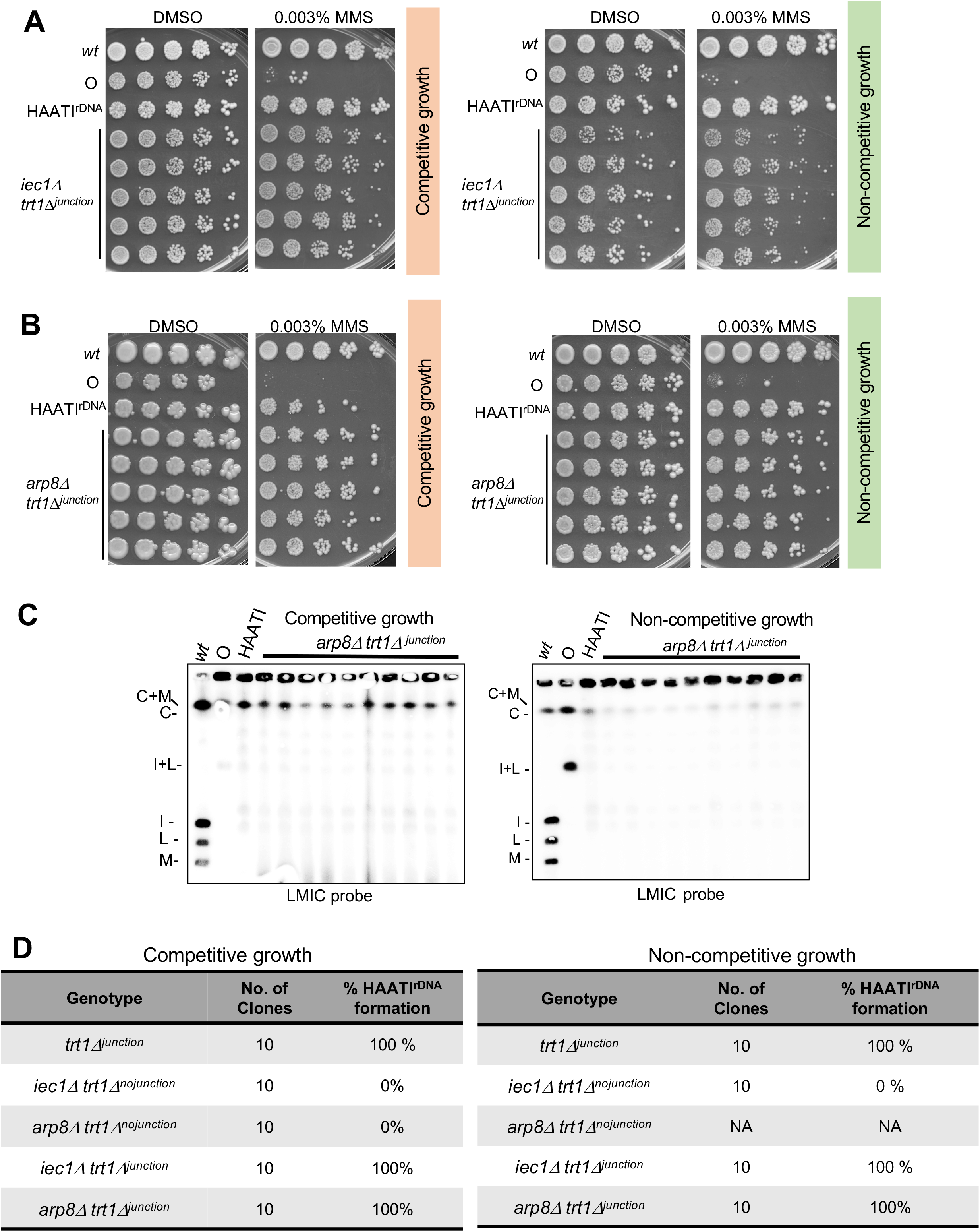
Ino80 complex is essential only for the first step of translocation. A. All *iec1Δ trt1Δ^unction^* survivors raised under competitive growth (left) or noncompetitive (right) conditions show MMS resistance suggestive of HAATI formation. B. All *arp8Δ trt1 Δi^unctton^* survivors propagated under either competitive (left) or noncompetitive conditions (right) show MMS resistance indicative of HAATI formation. C. PFGE of Not1 digested chromosomes of *arp8Δ trt1 Δi^unctton^* survivors. Retention of hybridization signal in the well for all survivors indicates HAATI formation. D. Table highlighting the dispensibility of Ino80C for HAATI^rDNA^ formation in the presence of a pre-existing STE-rDNA junction. Regardless of growth conditions, all *iec1Δ trt1Δ^unction^* or *arp8Δ trt1Δ^unctton^*survivors are HAATI. For comparison, *trt1 Δ?^unctton^* survivor formation data from Figures 3D and 6D is shown.

## Discussion

When telomeres erode, the chaotic fates that are continually prevented by functional telomeres become evident. Here we probe two such fates, the translocation of rDNA tracts from Chr III to nonhomologous sequences exposed by telomere loss on Chr I and II (to form HAATI^rDNA^), and the transposition-like mobilization of STE tracts from Chr I and II to multiple terminal and internal loci (to form HAATI^STE^). We find that HAATI^rDNA^ formation is sealed by a rare, initial translocation of rDNA to one STE chromosome end; this depends on the RNAi pathway and Ino80C. Once this single translocation has occurred, copying of the newly created STE-rDNA junction to other STE ends occurs easily and independently of RNAi and Ino80C. We also identify parallel roles for two individual players in these pathways, Dcr1 of the RNAi pathway and Iec1 of the Ino80C, that define noncanonical activities for these proteins in preventing HAATI^STE^ formation. Our results further delineate newly recognized translocation pathways whose impacts may extend beyond eroding telomeres.

### HAATI^rDNA^ formation

siRNAs and RNAi pathway components have been implicated in several facets of the DNA damage response (DDR) (31–35). In fission yeast, the siRNAs generated by RNAi target HC formation (31,36–38), which generally inhibits homologous recombination (HR) (39,40). Moreover, at pericentric regions where replication and transcription compete, co-transcriptional RNAi releases RNAPII, favoring completion of DNA replication and local HC propagation *via* replication-coupled histone modification, again preventing HR in the region (41,42). In the rDNA intergenic spacer region, Dcr1 has been specifically implicated in displacing RNAPII, which transcribes the intergenic spacer region in the direction opposing rDNA replication (12). Thus, Dcr1 resolves transcription-replication collisions, whose persistence in the absence of Dcr1 leads to elevated HR, again painting RNAi components as protectors of replication forks and inhibitors of recombination. A distinct set of roles for RNAi factors has been suggested in mammalian systems, in which transcription of regions adjacent to DSBs and damaged telomeres has been reported (33,35,43). The resulting ncRNAs are processed into siRNAs that are thought to serve as a platform or condensate that promotes the concentration of DDR factors (44).

As an ATP-dependent chromatin remodeler that both slides and exchanges nucleosomes, Ino80C has been implicated in a number of processes including DNA replication and repair (45–48). Ino80C shares with RNAi a role in RNAPII removal during replication stress, collaborating with transcription and proteasomal complexes to achieve RNAPII displacement (46,47). Moreover, Ino80C has also been shown to prevent spurious, pervasive transcription around replication origins (49). Ino80C has been shown to localize to R-loop enriched regions in the genome and counteract their formation (50). In addition, Ino80C has been implicated in reducing genomic nucleosome coverage and thereby mobilizing chromosomes in cells experiencing DSBs; this chromosome mobility promotes the homology search required for HR repair (46).

How might these foregoing observations inform our conception of why RNAi and Ino80C are absolutely required for the first illegitimate translocation that places rDNA on STE chromosome ends? One scenario entails that as telomere erosion abolishes TPE, enhanced STE and rDNA transcription at *trt1*Δ subtelomeres creates significant levels of transcription-replication collisions that threaten replication fork maintenance. Conceivably, replication forks need RNAi-based protection in order to create the substrate for a rare DNA polymerase template switch, from a stalled fork in the STE region to a stalled fork in the rDNA. Moreover, as repressive H3K9Me2/3-containing HC in the rDNA region is critical for preventing local R-loop accumulation (51), the loss of TPE as telomeres erode likely compromises this HC, necessitating Ino80C for RNAPII removal and R-loop clearance, both of which may be crucial for providing access to a DNA replication intermediate that serves as substrate for template switching. Ino80C-mediated reductions in nucleosome density (46,52) could also be crucial for allowing DNA- or RNA-polymerase template switching. Alternatively, or in addition, the accumulation of siRNAs may promote local condensate formation, which in turn may create a local hub that favors template switching reactions. Indeed, *in vitro* studies suggest that the binding of Ino80C to the general transcription factor-Taf14 promotes liquid-liquid phase separation (53). Thus, conceivably, the recruitment of Ino80C along with local enrichment of siRNAs may promote formation of a condensate that promotes translocation.

### Generating a genetic toolkit for HAATI^rDNA^ maintenance

Our work defines two types of HAATI regulators, those required solely for the initial translocation step leading to formation and those required for both formation and longterm maintenance of HAATI. Any factor required for HAATI maintenance will also be scored as required for formation, as maintenance over the duration of culturing is required for all assays of HAATI formation. For HAATI^rDNA^, the group of factors required solely for translocation comprises the RNAi pathway and Ino80C, while the maintenance group includes Reb1, Rad51, Pot1, Tpz1, Ccq1 and the HC assembly machinery. We speculate that the continued requirement for these factors reflects a scenario in which replication forks stall at the rDNA in a Reb1 dependent manner, triggering the generation of ss rDNA sequences that become coated with Rad51, which in turn collaborates with HC-associated Ccq1 to assemble Pot1-Tpz1 at rDNA chromosome ends and confer end-protection. This process needs to be repeated in every cell cycle to allow faithful propagation of HAATI^rDNA^ chromosome linearity.

### ‘Acceptor’ STE tracts: Specialized sequences or just ‘right place right time’?

HAATI^rDNA^ translocations occur between ‘donor’ rDNA sequences and ‘acceptor’ STE tracts, both repetitive subterminal HC regions. Like rDNA, STE tracts are poorly transcribed and subjected to TPE. STE transcription likely increases upon telomere erosion, creating potentially potent recombinogenic targets. Our observation that *STE*Δ strains fail to form HAATI^rDNA^ upon *trt1+* deletion suggests that STE sequences indeed harbor special HAATI^rDNA^-promoting characteristics. However, the interesting caveat that *STE* deletion shifts the adjoining knob regions to telomere adjacency appears to be crucial, as removal of the knob via *sgo2+* deletion allows for HAATI^rDNA^ formation, albeit at a low level. These data underscore the importance of chromatin structure in driving translocation choices; and the role of knob-mediated chromatin repression therein warrants further study.

### HAATI^STE^ formation

The mechanisms underlying both formation of, and end-protection by, HAATI^STE^ survivors remain exceedingly mysterious, with the observation that Dcr1 and Iec1 are crucial barriers to HAATI^STE^ formation constituting a rare hint. We speculate that STE transcripts whose production increases upon telomere erosion are normally cleaved by Dcr1, as the catalytic activity of Dcr1 is required for its blockage of HAATI^STE^ formation (while catalytic activity is dispensable for the role of Dcr1 in RNAPII eviction); in a *dcr1*Δ background, these transcripts would persist and integrate at multiple sites throughout the genome. Our attempts thus far to identify the junctions between STE and unique genomic sequences in HAATI^STE^ cells using whole genome sequencing have failed to yield such junctions, despite the use of long-read sequencing and high genomic coverage (unpublished data). Coupled with the persistence of STE hybridization to all the NotI fragments of the genome throughout the life of HAATI^STE^ cells, this observation suggests that STE sequences remain indefinitely mobile.

Iec1, a novel C2H2 zinc finger protein, co-purifies with Ino80, Rvb1/2 and nuclear actin to form a core Ino80C. This *S. pombe* core complex resembles the Yin-Yang1(YY1)/pleiohomeotic -Ino80 core complexes of higher eukaryotes with Iec1 as a *S.pombe* ortholog for YY1 (Hogan et al 2010). There is little precedent to allow us to speculate on how Iec1 blocks HAATI^STE^ formation in an Ino80C-independent manner, or on whether Iec1 blocks HAATI^STE^ translocations at the level of the ‘donor’ STE region or the ‘invaded’ recipient regions. Intriguingly, however, in the evolutionary distant invertebrate Lancelet, YY1 has been shown to suppress transposition of the ancestral RAG transposon ProtoRAG (54). Moreover, recombinant YY1 is itself sufficient to bind HR intermediates such as Holliday junctions and Y-shaped DNA intermediates (55). Conceivably, Iec1 shares this ability with YY1. By binding STE integration intermediates, perhaps Iec1 could promote their processing in a manner that prevents genome-wide erroneous integrations.

Given the clear blockage of HAATI^STE^ formation by Dcr1 and Iec1, the observation that *dcr1*Δ cells require Iec1 for HAATI^STE^ formation while *iec1*Δ cells require Dcr1 to form HAATI^STE^ is seemingly paradoxical. We suggest a scenario in which the simultaneous absence of Iec1 and Dcr1 lifts all the inhibitions that prevent aberrant STE ‘jumping’, triggering excessive levels of insertions across the genome, extreme genotoxic stress and cell death. In this scenario, the overzealous STE jumping would prevent HAATI^STE^ from dominating in *iec1Δ dcr1Δ trt1*Δ population, favoring circulars.

### Perspectives

Translocation of rDNA to nonhomologous sites is strictly confined to late generation *trt1*Δ cells whose telomeres have eroded. We speculate that evolution has placed rDNA in subtelomeric regions in order to prevent these translocations, and that similar forces have also restricted rDNA to the acrocentric chromosomes’ short arms in human cells as well. It will be interesting to determine whether any challenges to rDNA stability other than telomere loss might trigger such RNAi- and Ino80C-dependent translocations.

## Supplementary Data statement

Supplementary Data are available at NAR online.

## ACKNOWLEDGEMENTS

We thank all members of the JPC lab for their valuable comments and suggestions, with particular thanks to Rishi Kumar Nageshan for experimental help and advice. We thank the CCR sequencing facility at NCI Frederick for help with whole-genome sequencing; Karl Ekwall, Patrick Verga-Weisz and Junko Kanoh for sharing reagents and unpublished data; Michael Lichten and Eros Lazzerini Denchi for discussions and support. This work was supported by the National Cancer Institute and the University of Colorado School of Medicine.

## FUNDING

This work was supported by the National Institutes of Health and the University of Colorado School of Medicine.

Funding for open access charge: National Institutes of Health.

## CONFLICT OF INTEREST

The authors declare no conflict of interest.

## Supplementary Table and Figures

**Supplementary Table S1.**
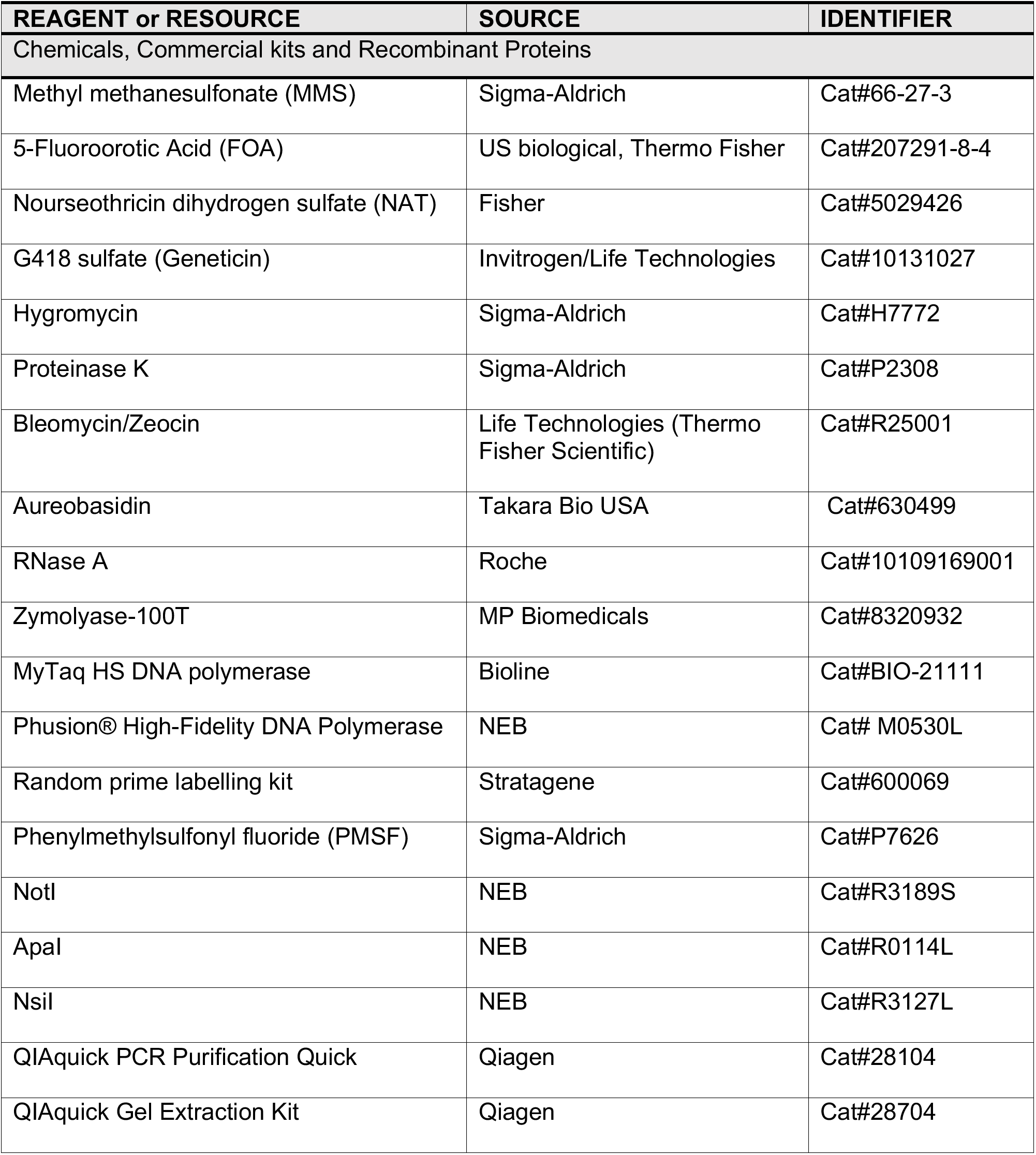
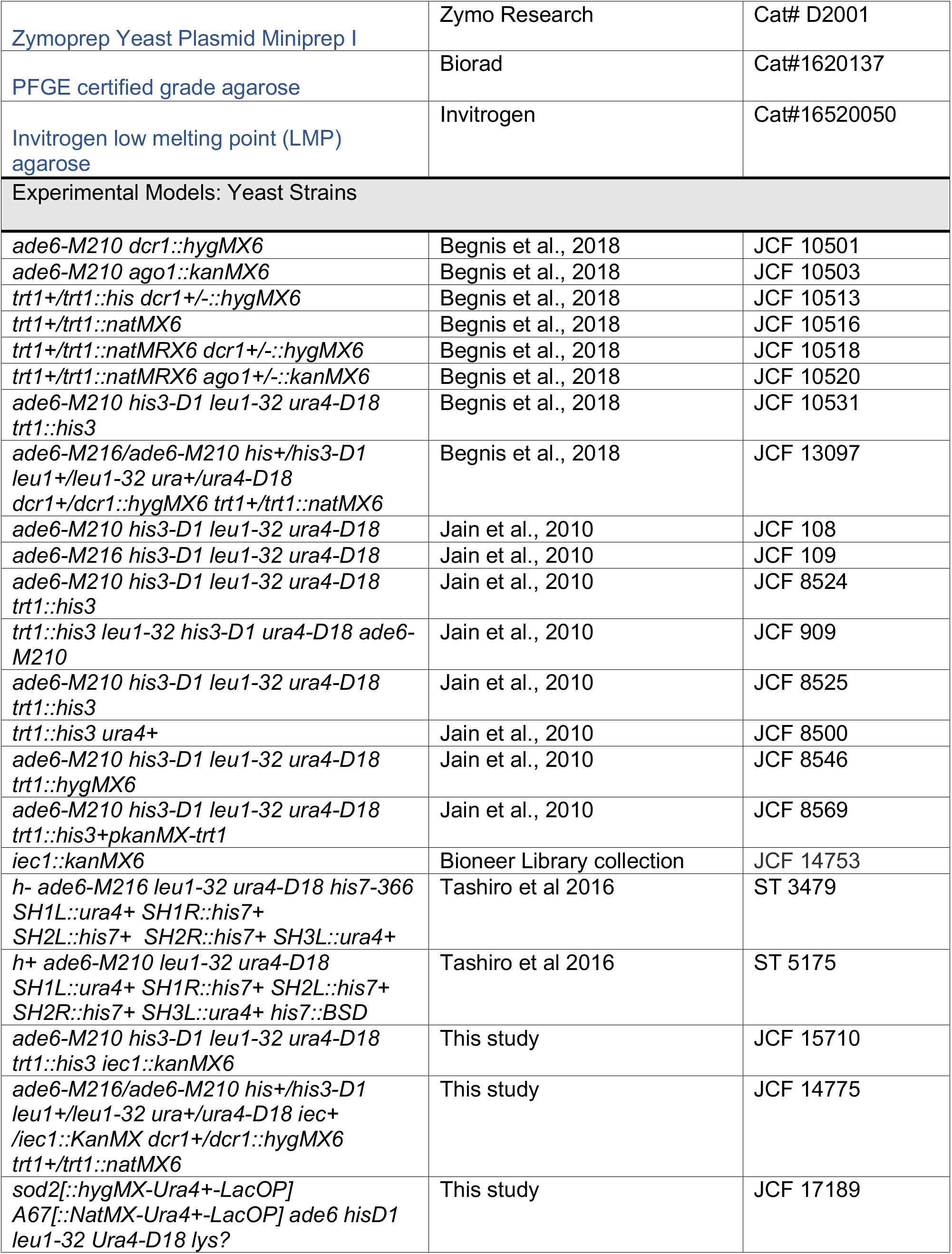
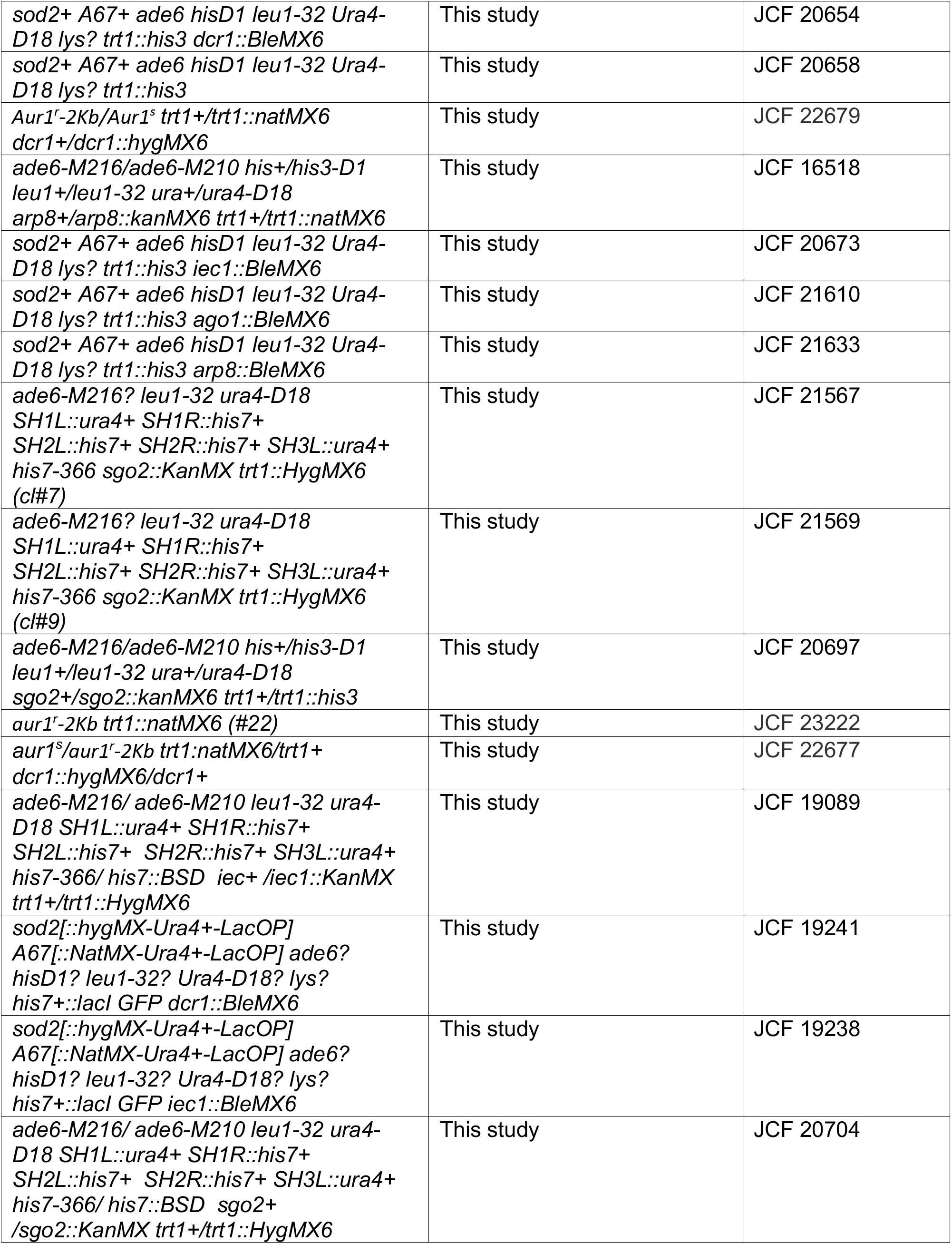
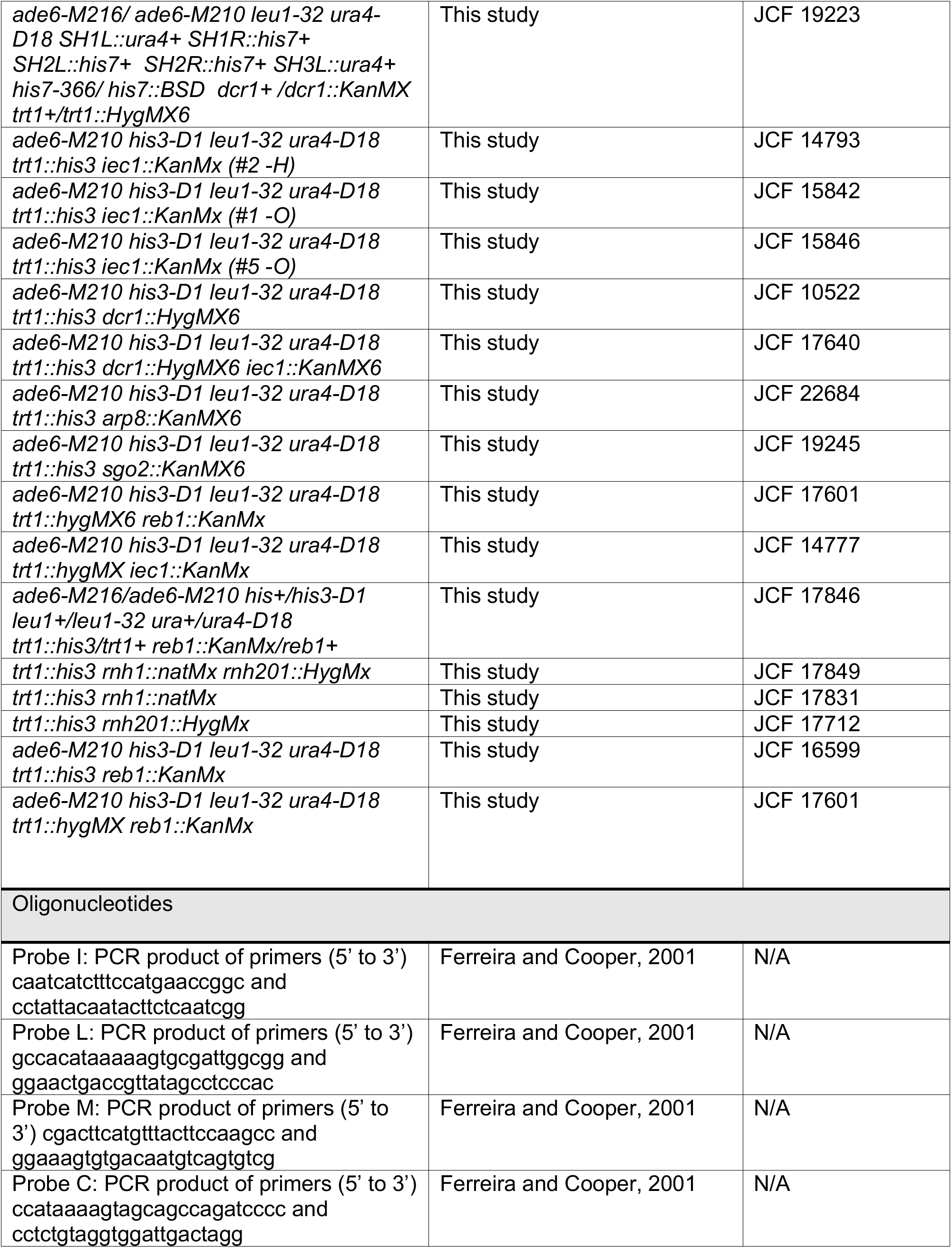
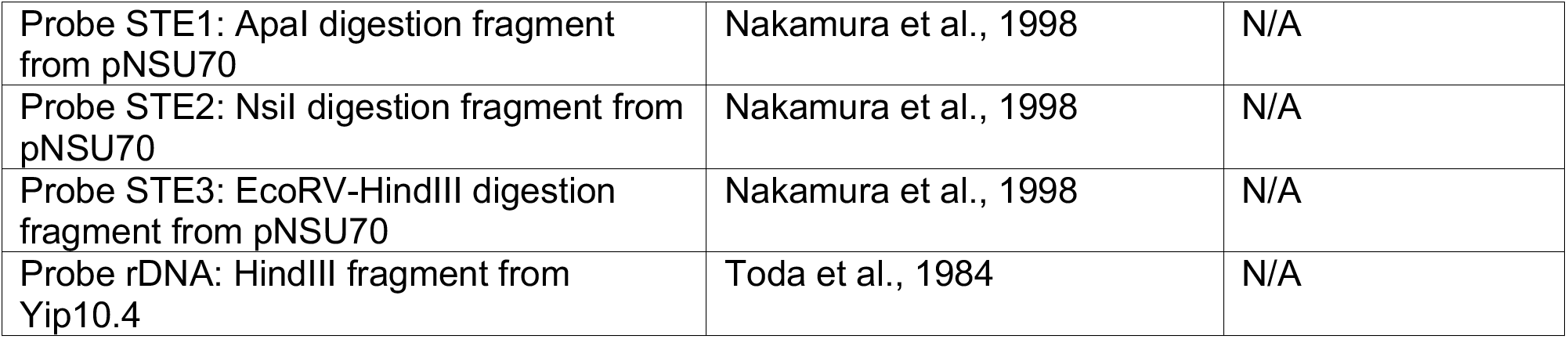
List of reagents, recombinant proteins, yeast strains and oligonucleotide primers is presented.

**Supplementary Figure S1:**
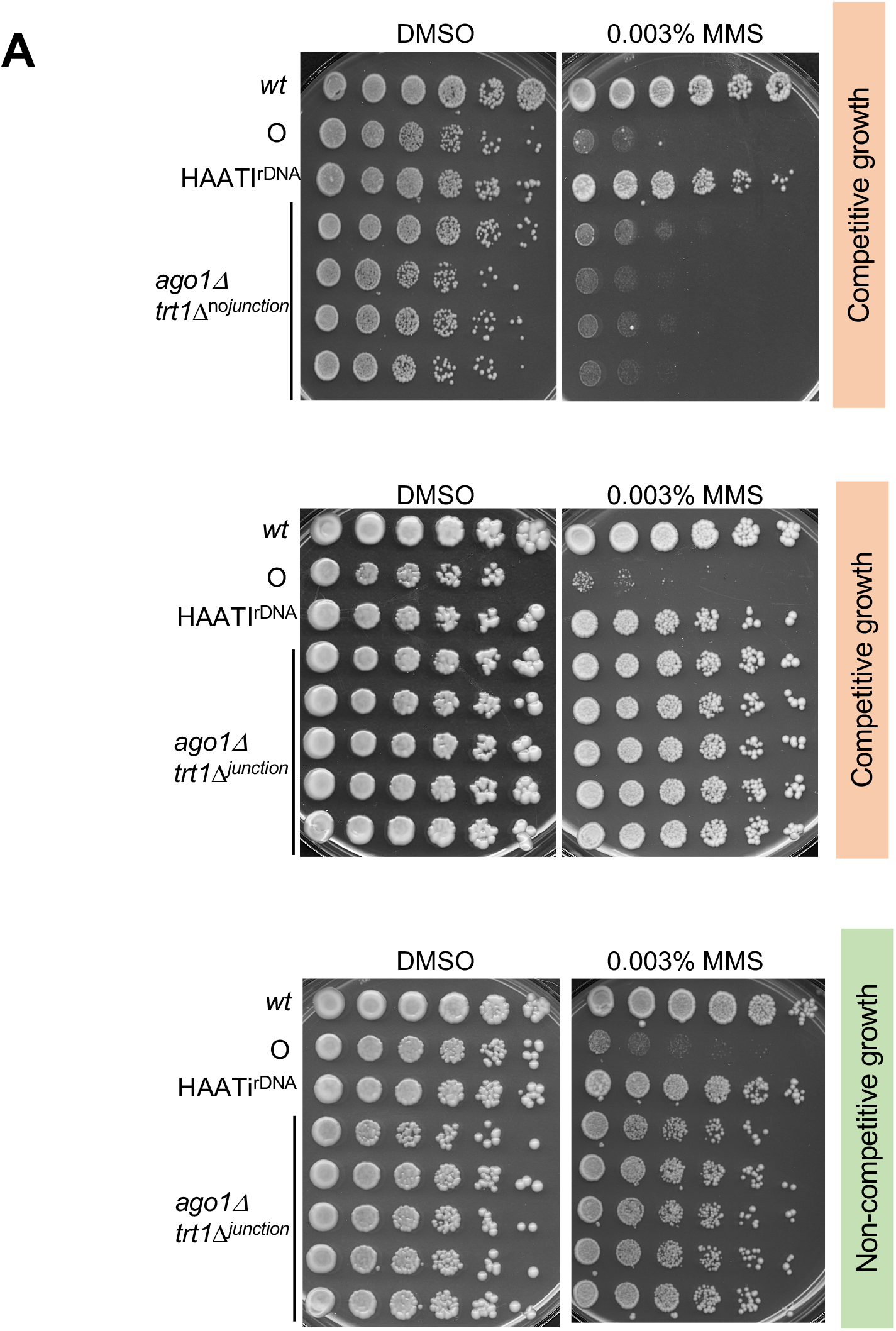
Ago1 is essential only for driving the first step of translocation. A. 5-fold serial dilutions of the indicated strains were spotted on media with or without MMS. (top) *ago1Δ trt1 Δ^nojunctton^* survivors grown under competitive conditions show MMS sensitivity suggesting chromosome circularization. In contrast, *ago1Δ trt1 Δ^unctton^* survivors propagated under either competitive (middle) or noncompetitive (bottom) growth conditions show MMS resistance indicative of HAATI formation.

**Supplementary Figure S2.**
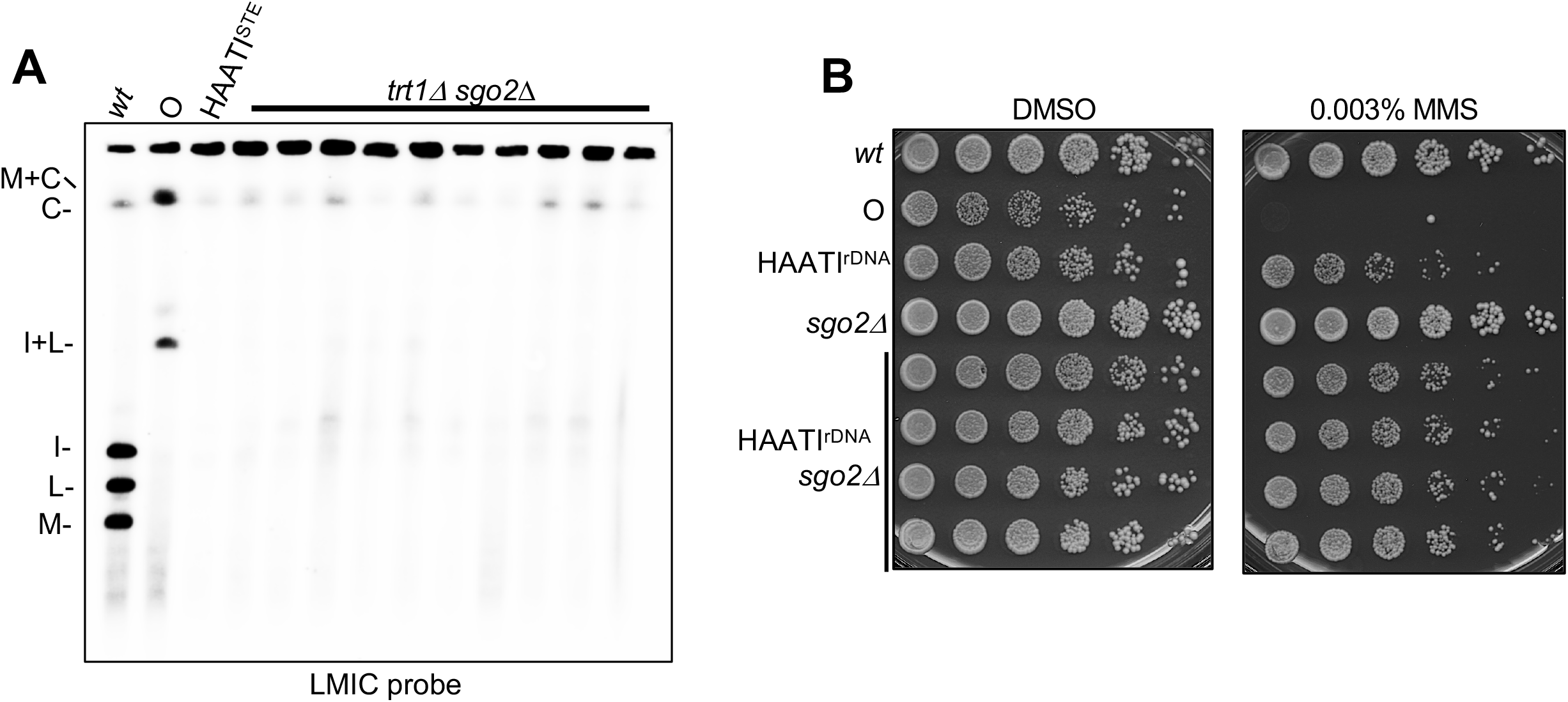
Sgo2 is dispensable for HAATI survival. A. PFGE of NotI digested chromosomes of *trt1Δ sgo2*Δ survivors. All survivors showed retention of LMIC signal in the wel, evincing HAATI formation. B. 5-fold serial dilutions spotted on media with or without MMS. All HAATI^rDNA^ clones retain MMS resistance upon loss of *sgo2Δ.*

**Supplementary Figure S3:**
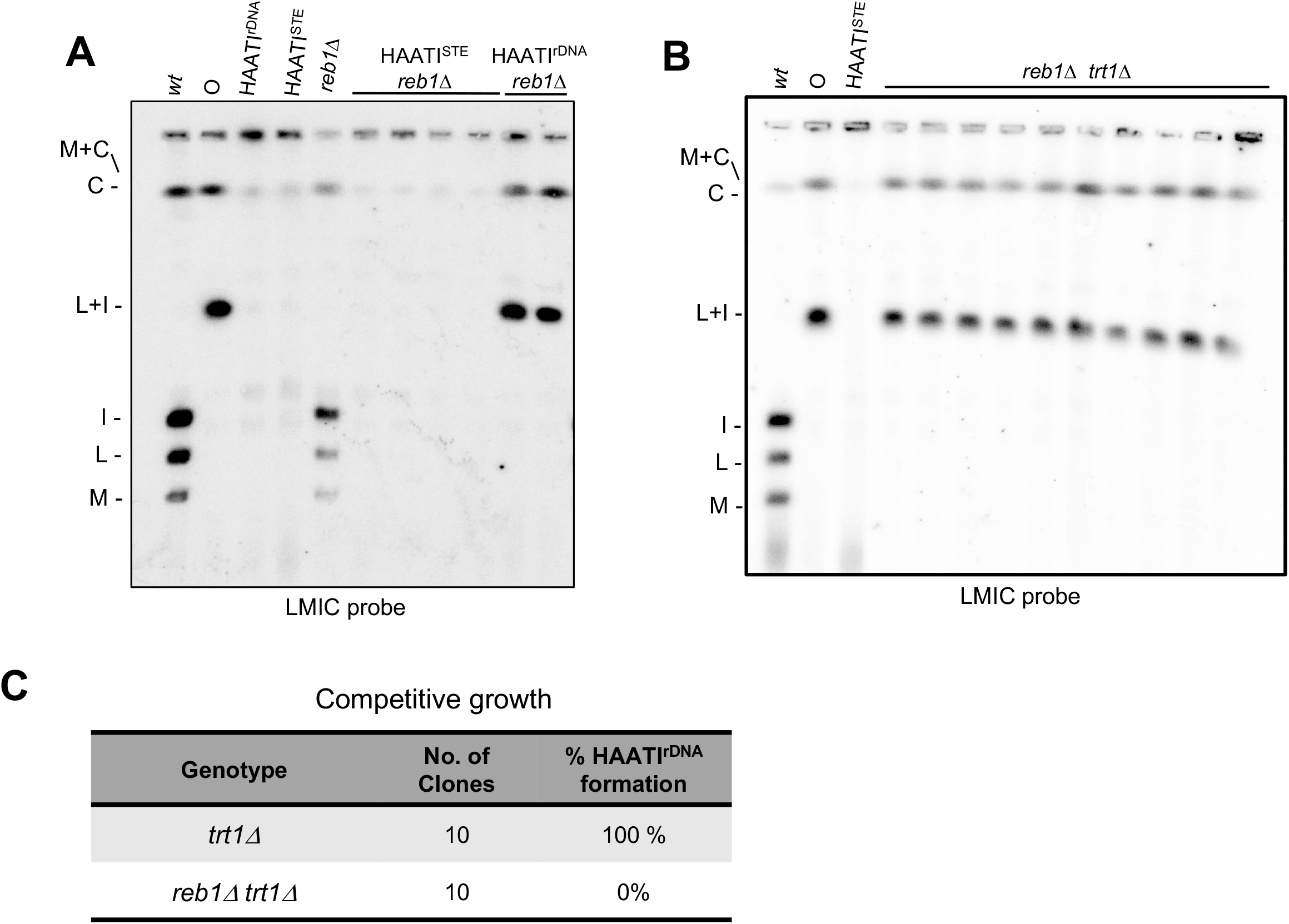
Reb1 is crucial for HAATI^rDNA^ survival. A. PFGE of NotI-digested chromosomes of HAATI^rDNA^ *reb1*Δ reveals bands evincing chromosome circularization while *reb1+* deletion fails to affect HAATI^STE^ pattern. B. PFGE analysis of progeny of heterozygous *trt1Δ/trt1+* diploids carrying or lacking Reb1. All *reb1Δ trt1*Δ survivors are circulars. C. Table summarizing the essential role of Reb1 in HAATI^rDNA^ formation. Percentage of HAATI^rDNA^ formation based on both MMS resistance and PFGE analysis. Under competitive growth conditions 10 out of 10 survivors of either *trt1*Δ formed HAATI but all survivors from *reb1Δ trt1*Δ were circular.

**Supplementary Figure S4.**
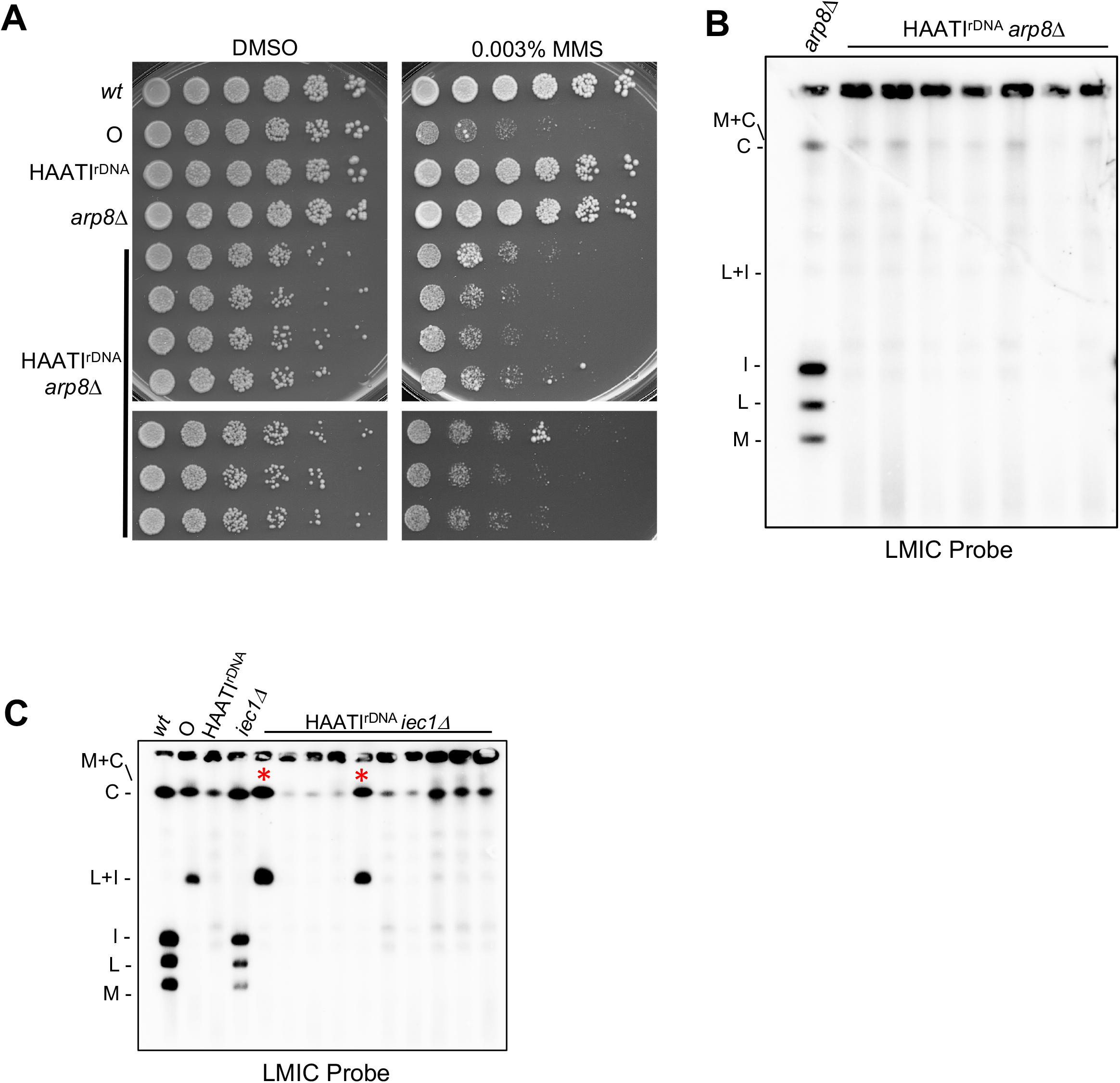
Ino80C is important but not essential for HAATI^rDNA^ chromosome linearity. A. 5-fold serial dilutions of the indicated strains were spotted on media with or without MMS. In pre-formed HAATI^rDNA^ cells, loss of *arp8^+^* yields slow growth and MMS sensitivity. B. PFGE of NotI-digested chromosomes from pre-formed HAATI^rDNA^ populations in which *arp8^+^* was deleted. All clones retain the HAATI^rDNA^ pattern. C. PFGE of NotI-digested chromosomes resulting from *iec1+*deletion in pre-formed HAATI^rDNA^ cells is shown. Two of 10 clones (asterisks) show chromosome circularization while others retain linear HAATI chromosome structure.

**Supplementary Figure S5:**
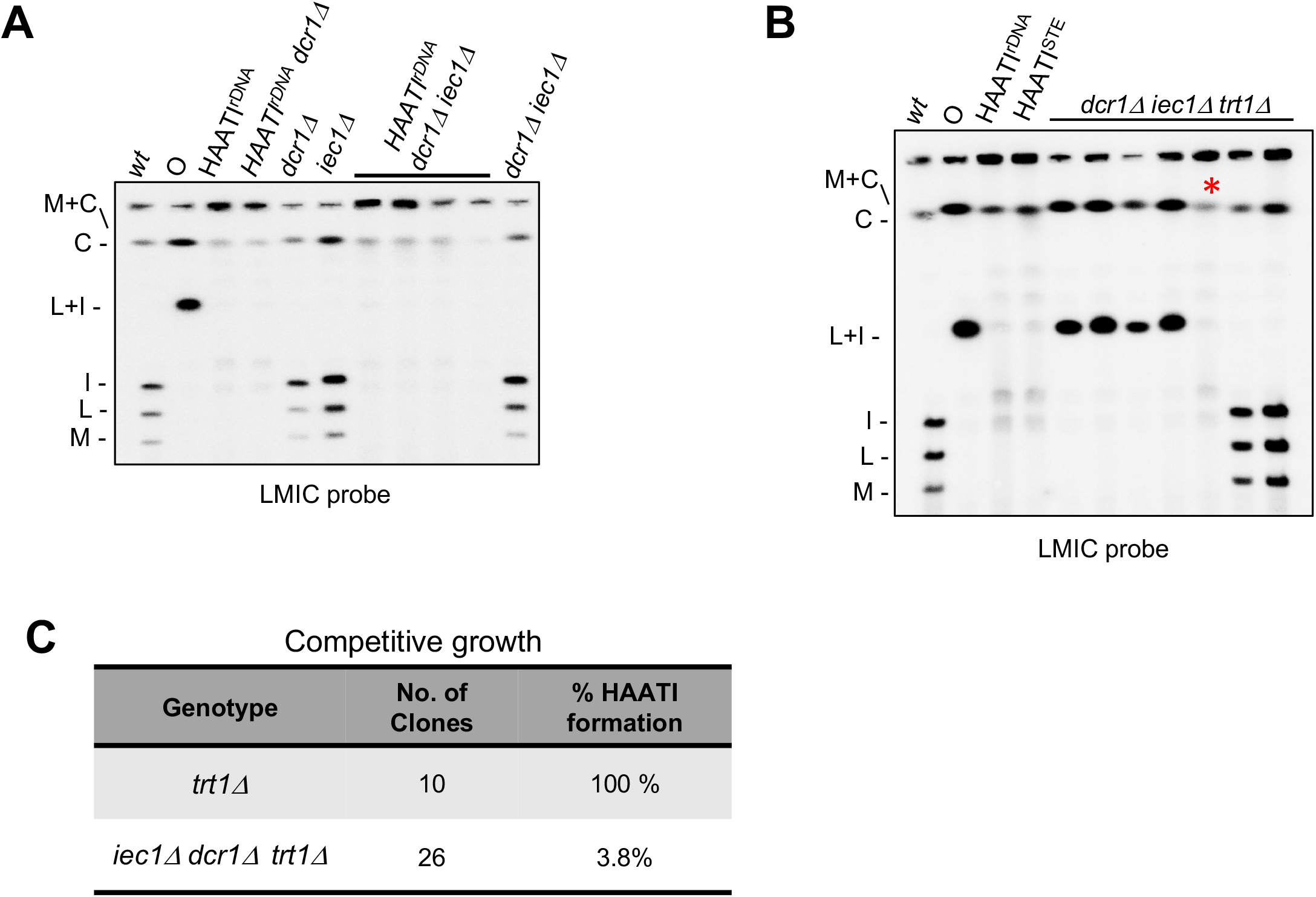
Iec1-Dcr1 crosstalk is required for HAATI formation. A. Representative PFGE of NotI-digested chromosomes from pre-formed HAATI^rDNA^ cells in which *dcr1+* and *iec1+* were deleted shows retention of linear HAATI^rDNA^ chromosome structure. B. PFGE analysis of progeny of heterozygous *trt1Δ/trt1+* diploids carrying or lacking both Iec1 and Dcr1. *iec1Δ dcr1Δ trt1*Δ fail to form HAATI. C. Table describing need for Dcr1 or Iec1 for HAATI^STE^ formation. Only one of 26 survivors show HAATI PFGE pattern..

## Notes

### Competing Interest Statement

The authors have declared no competing interest.

